# A fine-tuned genomic language model captures nucleotide-level information overlooked by missense variant impact predictors

**DOI:** 10.64898/2026.05.06.723362

**Authors:** Yaqi Su, Yu-Jen Lin

## Abstract

Established missense variant impact predictors primarily capture protein-level consequences, yet missense-annotated variants can also act through nucleotide sequence context. We developed GLM-Missense by systematically optimizing the fine-tuning of a genomic language model for missense variant impact predic-tion. GLM-Missense contributed the most complementary information to seven established predictors on held-out ClinVar variants and independent *RAD51D* saturation genome editing data. The complementary signal partly reflects splicing disruption, including disrupted exonic splicing regulatory elements largely missed by SpliceAI, and possibly codon-level effects on mRNA stability. These findings suggest fine-tuned genomic language models capture nucleotide-level information overlooked by current missense variant impact predictors.

## 1 Background

Missense variants are a major class of protein-altering coding variation, but their clinical interpretation remains challenging Majewski et al. (2011); Tennessen et al. (2012); McLaughlin et al. (2014). Clinical classification of missense variants is central to molecular diagnosis and variant prioritization. It requires consistent evaluation of evidence from diverse sources to ensure reproducible variant classification across lab-oratories. As next-generation sequencing brought large numbers of previously unseen variants into clinical testing, the American College of Medical Genetics and Genomics and the Association for Molecular Pathology (ACMG/AMP) developed joint guide-lines to standardize this process Richards et al. (2015). These guidelines introduced a common five-tier terminology (pathogenic, likely pathogenic, uncertain significance, likely benign, and benign) and a structured framework for weighing and combining evidence. Within this framework, the criteria PP3 and BP4 specifically address computational evidence supporting deleterious or benign effects, respectively. ClinVar aggregates clinical assertions of variant significance classified under this framework, including disease context and supporting evidence, and has become a key resource for variant interpretation Landrum et al. (2018, 2020). Nevertheless, many missense variants remain difficult to classify because clinical, segregation, and functional evidence are often limited Starita et al. (2017). This makes computational evidence particularly important for interpreting previously uncharacterized variants Pejaver et al. (2022).

Most existing missense variant impact predictors were designed to capture protein-level consequences, typically through amino-acid sequence conservation, structural context, and predicted effects on protein function Ng and Henikoff (2006). At the protein level, pathogenic missense variants can disrupt stability, catalytic activity, binding interfaces, or other biochemical functions, so these are natural targets for computational prediction Stefl et al. (2013). Current methods span several methodological classes. Early predictors such as SIFT and PolyPhen-2 estimate the functional impact of amino-acid substitutions from evolutionary conservation, alone or combined with physicochemical and structural properties Ng and Henikoff (2003); Adzhubei et al. (2010). Supervised predictors such as VEST4 and MutPred2 learn from larger sets of amino-acid substitution, conservation, and structural features Carter et al. (2013); Pejaver et al. (2020). Meta-predictors such as REVEL, BayesDel, and VARITY then combine the scores of multiple existing predictors with additional features Ioannidis et al. (2016); Feng (2017); Wu et al. (2021). More recently, protein language models including ESM1b and AlphaMissense, have demonstrated promising performance by leveraging large-scale proteomic and structural priors Brandes et al. (2023); Cheng et al. (2023). Large protein resources such as UniProt have enabled self-supervised pretraining, and protein language models have shown strong transfer across structural and functional tasks, including zero-shot mutation-effect prediction Bateman et al. (2023); Lin et al. (2023); Meier et al. (2021).

ClinGen has calibrated seven missense variant impact predictors (AlphaMissense, BayesDel, ESM1b, MutPred2, REVEL, VARITY, and VEST4) to provide computational evidence of defined strength under the ACMG/AMP PP3 and BP4 criteria Pejaver et al. (2022); Bergquist et al. (2025). Despite advances in missense variant impact prediction, independent benchmarking still reveals substantial residual error in distinguishing pathogenic from benign missense variants Rastogi et al. (2025). It remains unclear whether these remaining errors mainly reflect irreducible uncertainty or biological information overlooked by existing predictors.

A missense annotation describes the encoded amino-acid substitution, but the underlying nucleotide change may also act through nucleotide sequence context. For example, exonic variants can alter splicing by perturbing splice-adjacent sequence, exonic splicing enhancers or silencers, or cryptic splice sites Fairbrother et al. (2002); Cartegni et al. (2002); Jaganathan et al. (2019). Previous studies estimated that approximately 10% of disease-causing exonic mutations alter splicing, and missense exonic disease mutations with the least predicted protein-level impact are especially enriched for splicing disruption Soemedi et al. (2017); Xiong et al. (2015). Coding DNA also carries information beyond the encoded amino-acid sequence. Codon usage can influence mRNA stability and translation Presnyak et al. (2015); Wu et al. (2019). Local nucleotide changes can perturb RNA structure Sabarinathan et al. (2013). Coding exons can contain overlapping regulatory code, including dual-use codons that also encode transcription-factor-binding information Stergachis et al. (2013). The mis-sense annotation can therefore obscure a broader set of mechanisms than protein-level consequence alone. We therefore hypothesized that part of the remaining error in cur-rent missense variant impact prediction arises because many existing methods focus primarily on protein-level effects and do not fully capture nucleotide-level context.

Genomic language models offer a way to address this gap by learning representations directly from DNA sequence. DNABERT first showed that self-supervised learning on genomic sequence can produce transferable representations for down-stream genomic tasks Ji et al. (2021). Subsequent models have explored different tokenization schemes, model scales, pretraining corpora, and architectures, ranging from Transformer-based models such as the Nucleotide Transformer family Dalla-Torre et al. (2025) and DNABERT-2 Zhou et al. (2024) to long-context models based on implicit convolutions or state-space layers, such as HyenaDNA and Caduceus Nguyen et al. (2023); Schiff et al. (2024). Independent benchmarks have evaluated how these architectural and pretraining differences translate into downstream performance. BEND found that frozen embeddings from genomic language models approach expert methods on some tasks but capture limited long-range information Marin et al. (2024). Feng et al. and Genomic Touchstone further showed that model rankings vary substantially across genomic and genetic tasks Feng et al. (2025); Wang et al. (2025). Multi-species Nucleotide Transformer models, including Nucleotide Transformer v2 (NT-2), performed well on several variant effect tasks, including distinguishing disease-associated from benign or common variants Marin et al. (2024); Feng et al. (2025); Wang et al. (2025). State-space models such as Caduceus performed well on regulatory-element classification and splice variant effect prediction, but generally underperformed Transformer-based models on tasks requiring precise token-level predictions Feng et al. (2025); Wang et al. (2025). Together, these benchmarks show that model performance is task-dependent, motivating backbone selection based on the intended application. However, they primarily compare models under standardized evaluation protocols rather than systematically optimizing adaptation for a specific task.

Our study builds on these benchmarks by focusing on a single task: clinical missense variant impact prediction. We asked two questions. First, how should a pretrained genomic language model be adapted for missense variant impact prediction, given that labeled variants are limited relative to model size? Second, can the adapted model capture nucleotide-level information overlooked by established protein-centric missense variant impact predictors? To address these questions, we explored the fine-tuning strategies using NT-2, a multi-species Nucleotide Transformer v2 model, as our primary backbone Dalla-Torre et al. (2025). We chose NT-2 because it performed well on variant effect prediction tasks in the previous benchmarks Marin et al. (2024); Feng et al. (2025); Wang et al. (2025). We also evaluated two alternative back-bones. NT-1, a first-version Nucleotide Transformer model pretrained only on human genomes Dalla-Torre et al. (2025), allowed us to examine the potential contribution of multi-species pretraining. Caduceus-PS, a state-space model with built-in reverse-complement symmetry Schiff et al. (2024), allowed us to compare different model architectures.

Task-specific adaptation is important for missense variant impact prediction, since zero-shot scores from genomic language models have shown limited performance on this task Benegas et al. (2025); Curtis (2025). Choosing an effective adaptation strategy is critical because the number of clinically labeled missense variants are small relative to the size of current pretrained genomic models, and thus downstream performance may depend strongly on how the pretrained backbone is adapted Veiner and Supek (2026); Feng et al. (2025). Key design choices include how backbone hid-den states are aggregated into variant representations, which classifier head is used, how the reference and alternate alleles are encoded, and whether the backbone is kept frozen, fully fine-tuned, or adapted with parameter-efficient methods such as low-rank adaptation (LoRA) Hu et al. (2022). LoRA keeps the pretrained weights fixed and learns only small low-rank updates; by limiting the number of trainable parameters, it can help reduce overfitting when labeled data are limited relative to model size.

Ensemble models provide a framework for evaluating whether genomic language models capture information overlooked by established predictors. By combining scores from multiple predictors, ensembles can integrate partially complementary information and improve predictive performance. Meta-predictors such as REVEL demonstrate the benefits of this approach for missense variant impact prediction Ioannidis et al. (2016). Such improvements depend on the extent to which individual predictors con-tribute complementary information Dizkirici Tekpinar and Tekpinar (2025). Beyond aggregating signals, ensembles can also be used to quantify the additional information contributed by a new predictor. However, most benchmarking studies evaluate only standalone performance, which does not capture this contribution. We there-fore use ensemble ablation analysis to test whether a fine-tuned genomic language model captures information missing from established missense variant impact predictors. Specifically, we compare ensemble performance with and without the genomic language model while keeping the other predictors fixed. A performance decrease upon its removal indicates that it contributes predictive information not fully recovered by the remaining predictors.

Variant effect prediction benchmarks are vulnerable to circularity and other sources of optimistic bias Grimm et al. (2015); Livesey and Marsh (2020, 2023); Rastogi et al. (2025). Chromosome-level holdouts reduce potential data leakage by ensuring that variants from the same chromosome do not appear in both training and validation sets Whalen et al. (2022); Feng et al. (2025). However, clinical labels may introduce another source of circularity. Computational predictions may contribute to ClinVar classifications through the PP3 and BP4 criteria Pejaver et al. (2022). Evaluating the same predictors against these classifications can inflate apparent accuracy because their predictions may have helped determine the benchmark labels. Multiplexed assays of variant effect provide an alternative source of evaluation datasets by experimentally measuring the functional consequences of variants, independent of computational pre-dictions Fowler and Fields (2014); Weile and Roth (2018). Although variant function is only a proxy for pathogenicity, functional classifications from well-validated assays agree closely with clinical classifications Findlay et al. (2018). Saturation genome edit-ing (SGE), in particular, introduces nearly all possible single-nucleotide variants at the endogenous genomic locus, so that its functional readout reflects the consequence of each nucleotide change in its native sequence context, including effects at the RNA level, such as aberrant splicing, in addition to effects on the encoded protein Findlay et al. (2014, 2018).

Here, we systematically fine-tuned genomic language models to show that they can capture nucleotide-level information overlooked by established missense variant impact predictors (Fig. 1). Using NT-2 as the primary backbone, with NT-1 and Caduceus-PS as comparisons, we examined how pretraining corpus, embedding aggregation strategy, classifier head, and adaptation regime influence downstream performance. Variant-position embeddings outperformed mean-pooled sequence representations, multi-species pretraining provided the clearest backbone-level advantage, and LoRA generalized better than full fine-tuning under limited data. Building on the best-performing configuration, we developed GLM-Missense, a genomic language model fine-tuned for missense variant impact prediction. GLM-Missense substantially improved over zero-shot scoring from the same pretrained backbone. We then combined GLM-Missense with the seven comparator missense variant impact predictors calibrated by ClinGen for clinical use Bergquist et al. (2025) in MetaMissense, an XGBoost-based ensemble model, which outperformed each of the eight predictors on held-out ClinVar variants. Among the input predictors to MetaMissense, GLM-Missense showed the lowest concordance with the others and retained the strongest unique predictive signal. To quantify each predictor’s contribution to ensemble performance, we compared ensembles trained identically with and without that predictor. Removing GLM-Missense caused the largest loss of the ensemble’s ability to dis-criminate pathogenic from benign variants, on both held-out ClinVar variants and on an independent *RAD51D* SGE dataset Casadei et al. (2026). A protein language model fine-tuned on the same training variants did not reproduce this contribution, suggesting that the unique contribution of GLM-Missense arises from the nucleotide-level information it captures rather than from training on recent ClinVar labels alone. Among variants correctly classified by GLM-Missense but misclassified by most established predictors, we identified splicing disruption as one component of GLM-Missense’s distinctive signal, including disruption of exonic splicing regulatory elements that was missed by SpliceAI, a state-of-the-art splicing predictor Jaganathan et al. (2019). Finally, an analysis of variants of uncertain significance in ClinVar suggested that GLM-Missense may also capture codon-level effects on mRNA stability.

**Fig. 1.**
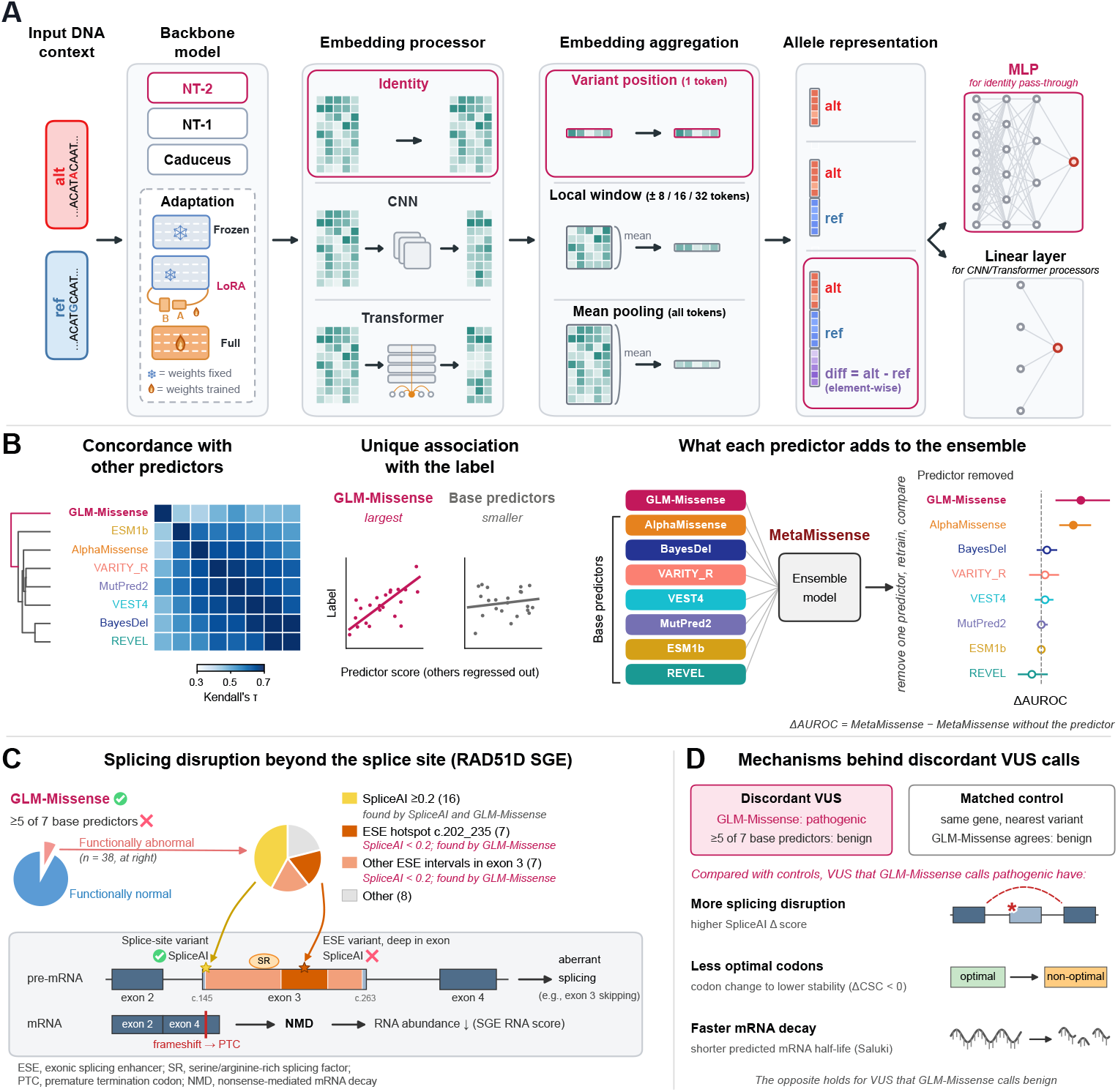
Schematic overview of the study. **(A)** Design space explored for adapting genomic language models to clinical missense variant impact prediction. We compared three genomic language model backbones (NT-2, NT-1 and Caduceus-PS), three adaptation regimes (frozen backbone, LoRA fine-tuning and full fine-tuning), three classifier or representation heads (identity, CNN and shallow Transformer), multiple embedding aggregation strategies (variant-position embedding, local-window mean pooling and full-sequence mean pooling), and three allele-aware representations (*alt*, *ref-alt* and *ref-alt-diff*). The best-performing configuration colored in raspberry pink was used to construct GLM-Missense. **(B)** GLM-Missense contributes information beyond the seven comparator missense variant impact predictors (BayesDel, MutPred2, REVEL, VEST4, AlphaMissense, ESM1b and VARITY_R). It showed the lowest concordance with these predictors, retained the strongest unique association with the label after accounting for them, and caused the largest drop in ensemble AUROC when removed from the MetaMissense ensemble on held-out ClinVar variants. **(C)** Splicing-related effects associated with missense variants in the *RAD51D* SGE dataset resolved by GLM-Missense (classified correctly by GLM-Missense but misclassified by most established predictors). These variants showed evidence of splicing disruption, including disruption of exonic splicing enhancers away from canonical splice sites that is overlooked by SpliceAI. **(D)** Variants of uncertain significance on which GLM-Missense disagrees with most established predictors also differ from matched controls in codon optimality and predicted mRNA half-life.

## 2 Results

### 2.1 Variant-position embeddings and multi-species pretraining yield the best frozen-backbone configuration

We first asked which frozen-backbone configuration provided the strongest baseline for clinical missense variant impact prediction. In this setting, the backbone parameters were fixed, and only a lightweight downstream classifier was trained. All frozen-backbone experiments used the ClinVar earlier-release strict set, defined as variants present in the November 3, 2025 ClinVar release with Benign and Pathogenic variants only, excluding likely and uncertain classifications (Methods). We compared three pretrained genomic language model backbones: NT-2, NT-1, and Caduceus-PS Dalla-Torre et al. (2025); Schiff et al. (2024). For each backbone, we evaluated multiple combinations of embedding aggregation strategies and classifier heads to identify the optimal baseline configuration (Fig. S1).

Embedding choice had a significant effect on performance. We first compared the embedding of the token containing the variant, which we call the *variant-position embedding*, against mean pooling across the full input sequence. Each representation was paired with an MLP, a CNN, or a shallow Transformer head. For Caduceus-PS, we first performed downsampling of the full sequence to account for memory constraints when paired with CNN or Transformer head. Variant-position embeddings consistently outperformed mean pooling in the models tested (Fig. 2A, Fig. S2).

**Fig. 2.**
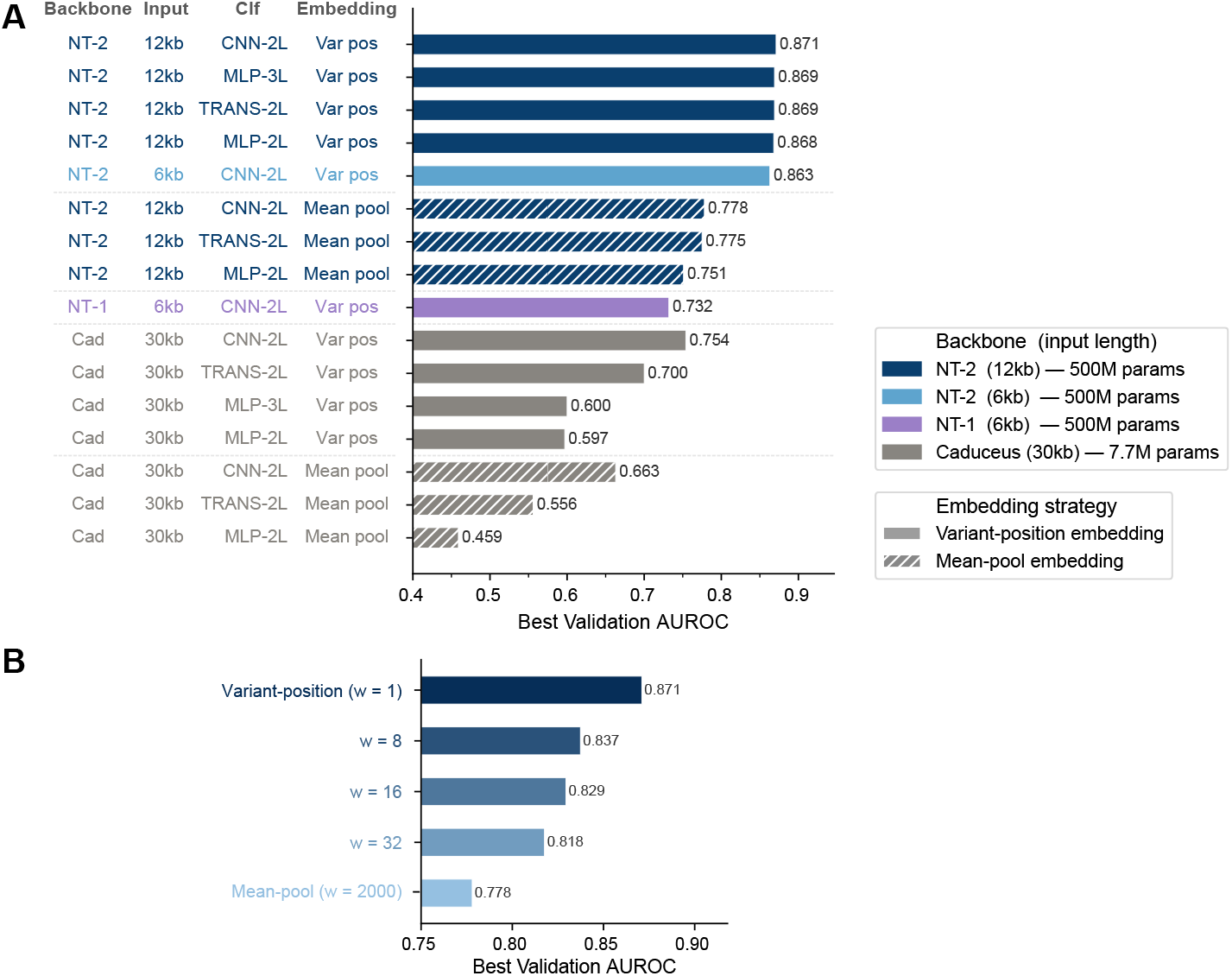
Frozen-backbone comparison across genomic language model backbones, embed-ding strategies, classifier heads, and window pooling sizes. **(A)** Best validation AUROC is shown for combinations of three backbone architectures: Nucleotide Transformer v2 (NT-2, 500M parameters), Nucleotide Transformer v1 (NT-1, 500M parameters), and Caduceus-PS (Cad, 7.7M parameters), with four classifier (Clf) heads: two-layer CNN (CNN-2L), three-layer MLP (MLP-3L), two-layer MLP (MLP-2L), and two-layer Transformer (TRANS-2L). Models were evaluated using either a variant-position embedding, which extracts the representation at the variant site, or a mean-pool embedding, which averages representations across the input sequence. Bar color denotes backbone and input length, while hatching denotes embedding strategy. **(B)** Effect of pooling window size around the variant position for the best-performing NT-2 frozen configuration. The variant-position embedding with window size w = 1 achieved the highest validation AUROC, while progressively larger pooling windows reduced performance.

We next examined whether pooling embeddings within a local window around the variant could preserve local context without averaging over irrelevant positions. To test this, in the frozen NT-2 model with a CNN classifier head, we replaced full-sequence mean pooling with mean pooling over ±8, ±16, or ±32-token windows centered on the variant (Fig. 2B, Fig. S3). Performance decreased progressively as the window widened, indicating that representation at the variant position was more informative than averaging across nearby tokens.

Classifier-head choice had a smaller effect than embedding choice. In NT-2 with a variant-position embedding, the three classifier heads achieved similar validation AUROCs of 0.869–0.871 (Fig. 2A, Fig. S2). In NT-2 with mean pooling, the spread across heads increased to ΔAUROC = 0.027. This pattern suggests that when suboptimal input representations are used, more sophisticated classifier architectures, such as CNN and Transformer, may partially compensate by learning to extract relevant features from the aggregated sequence information. The Caduceus-PS model consistently underperformed NT-2 in the frozen backbone setting (Fig. 2A, Fig. S2). Also noticeably, the performance among different classifier heads under the same embedding strategy varies more significantly for Caduceus-PS compared to NT-2, likely due to the weaker embedding representation generated by the Caduceus-PS backbone. Due to GPU memory constraints, the Caduceus-PS hidden states were downsampled before being passed to the CNN and Transformer heads, potentially reducing their performance. However, Caduceus-PS with the MLP configurations, which used the variant-position embedding directly without downsampling, still substantially under-performed NT-2 under otherwise identical settings (validation AUROC 0.597–0.600 compared with 0.868–0.869; Fig. 2A). This finding is consistent with an independent benchmark reporting that state-space genomic language models often underperformed Transformer-based models on tasks that require precise token-level predictions Wang et al. (2025).

The NT-2 backbone pretrained on multi-species genomes substantially outperformed the NT-1 backbone pretrained only on human genomes. To assess backbone performance under matched experimental conditions, we applied the same downstream configuration to both models and matched their input lengths at 6kb. NT-2 achieved a validation AUROC of 0.863, whereas NT-1 achieved 0.732 (Fig. 2A). Several factors may contribute to this performance difference, as NT-2 and NT-1 differ in their pre-training corpora, architectures, and pretraining hyperparameters. The contribution of pretraining corpus was examined more directly in the Genomic Touchstone bench-mark, which compared two Nucleotide Transformer v1 models that differed only in their pretraining corpora, while sharing the same architecture and model size (NT-2.5B-1000G and NT-2.5B-Multi). The multi-species model generally outperformed the model pretrained on human genomes only Wang et al. (2025). Together, these results suggest that multi-species pretraining likely contributes to the performance advantage of NT-2 in our task.

Overall, the frozen-backbone experiments identify two main design principles: pre-serving the representation at the variant position, and using a multi-species pretrained backbone. We therefore selected NT-2 as the backbone for subsequent adaptation experiments.

### 2.2 LoRA preserves variant-position advantages, improves generalization, and benefits from reference–alternate difference representations

We next asked whether the design principles from the frozen setting persisted under LoRA, where the pretrained backbone was allowed to adapt only through a small set of low-rank parameters Hu et al. (2022). The LoRA rank, denoted *r*, defines the dimension of these update matrices and therefore the number of trainable adaptation parameters. We evaluated 54 LoRA configurations on NT-2. The sweep varied embedding strategy (variant-position and mean pooling), classifier head (MLP, CNN, Transformer), LoRA rank (*r* ∈ {8, 16, 32}), and learning rate (1 × 10*^−^*^5^, 3 × 10*^−^*^5^, 5 × 10*^−^*^5^). To enable rapid iteration, these experiments used a reduced 5k-variant subset from the ClinVar earlier-release strict training set (Fig. 3A).

**Fig. 3.**
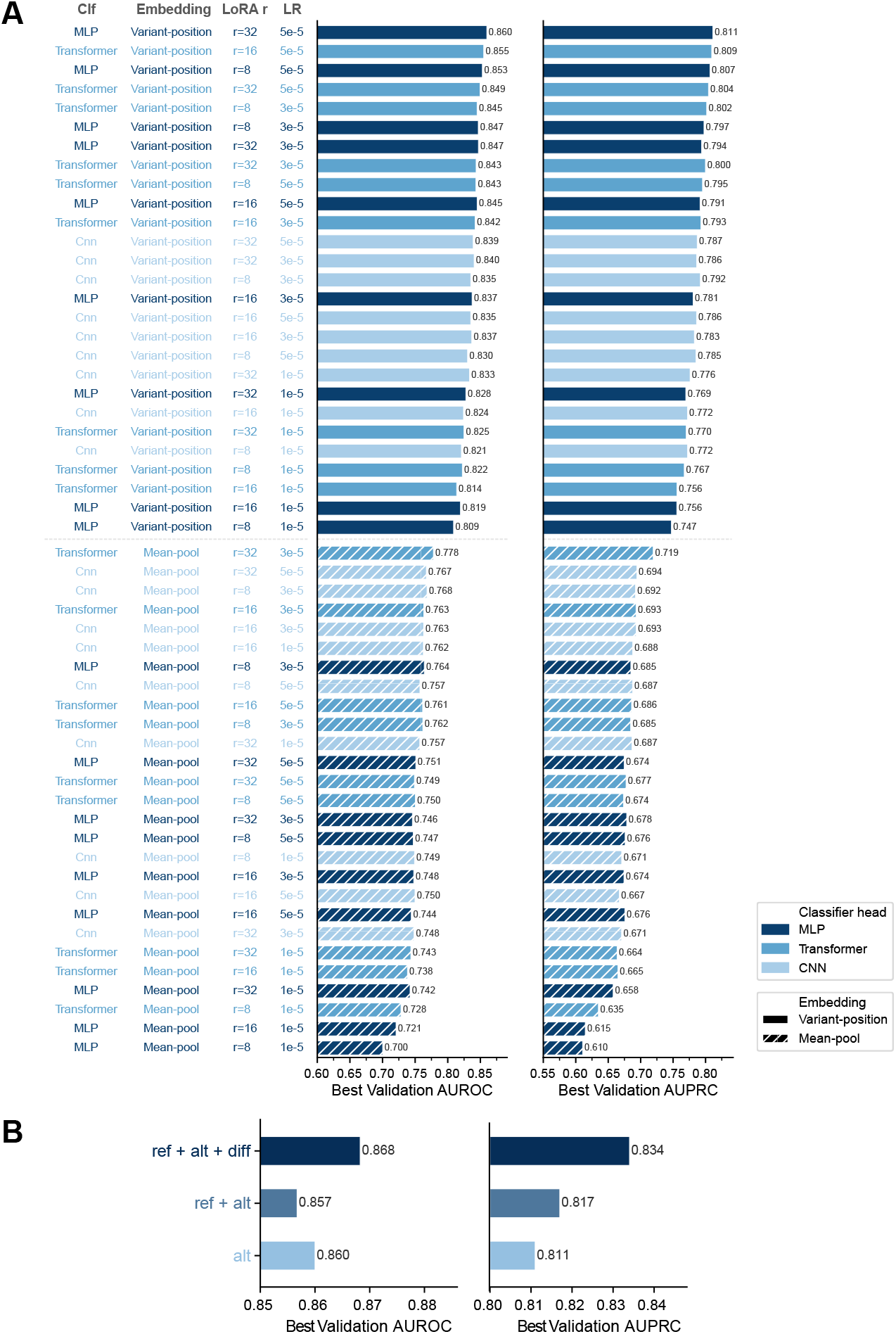
LoRA fine-tuning performance across classifier heads, embedding strategies, LoRA ranks, learning rates, and input feature representations. **(A)** Validation AUROC and AUPRC for 54 LoRA fine-tuning configurations. Each row corresponds to one combination of classifier head, embedding strategy, LoRA rank, and learning rate. Configurations are ordered by their mean rank across AUROC and AUPRC. Bar color denotes classifier head: MLP, Transformer, or CNN. Solid bars indicate variant-position embeddings, while hatched bars indicate mean-pool embeddings. **(B)** Exploration of input feature representations using the best-performing LoRA fine-tuning setup. ref + alt + diff uses the concatenation of reference allele embedding, alternate allele embedding, and their difference vector; ref + alt uses the concatenation of reference and alternate allele embeddings; and alt uses only the alternate allele embedding.

Variant-position embeddings remained the dominant design choice under LoRA. All 27 variant-position configurations outperformed all 27 mean-pooling configurations. The worst variant-position model (MLP, *r* = 8, lr = 1 × 10*^−^*^5^) achieved an AUROC of 0.809 and an AUPRC of 0.747. The best mean-pooling model (Trans-former, *r* = 32, lr = 3 × 10*^−^*^5^) reached only 0.778 AUROC and 0.719 AUPRC. Thus, even when the backbone was allowed to adapt through LoRA, preserving the representation at the variant position remained more effective than averaging across the sequence.

The MLP classifier performed best overall among LoRA models with variant-position embeddings (Fig. 3A). The best CNN configuration reached an AUROC of 0.840, compared with 0.860 for the top MLP configuration. This pattern suggests that once NT-2 is adapted through LoRA, an MLP head is sufficient to use the variant-position representation effectively. LoRA rank and learning rate had comparatively modest effects once the embedding strategy was fixed, with the top configurations closely clustered. To confirm that model selection was not driven by a single validation split, we evaluated the top ten LoRA settings by 5-fold cross-validation (Supplementary Table S1). The same MLP, variant-position configuration with *r* = 32 and lr = 5 × 10*^−^*^5^ remained the top-ranked setting after cross-validation, and was therefore selected for subsequent experiments.

We then scaled the selected LoRA configuration to the full 44k ClinVar earlier-release strict training set and compared it with full fine-tuning of NT-2 across five learning rates from 1 × 10*^−^*^6^ to 1 × 10*^−^*^4^ (Fig. S4). All runs used the same early stopping rule: training stopped when the train–validation gap exceeded 0.1 or when the validation metric failed to improve for 2,000 consecutive steps. Four full fine-tuning runs (1 × 10*^−^*^6^ to 3 × 10*^−^*^5^) stopped because the train–validation gap crossed the 0.1 threshold. In these runs, training AUROC and AUPRC continued to rise while validation performance plateaued or declined, indicating patterns of overfitting. The most aggressive full fine-tuning learning rate (1 × 10*^−^*^4^) stopped for failure to improve and remained near chance throughout training. By contrast, the LoRA run stopped only after validation performance plateaued at a higher level while maintaining a smaller train–validation gap.

Finally, we evaluated whether explicitly encoding the reference-to-alternate change improves predictive performance (Fig. 3B). The baseline *alt* representation used only the alternate allele variant-position embedding. The *ref-alt* representation concate-nated the reference and alternate embeddings, whereas *ref-alt-diff* concatenated the reference embedding, alternate embedding, and their element-wise difference. The *ref-alt-diff* representation performed the best, improving over the *alt* baseline by 0.008 AUROC and 0.023 AUPRC. In contrast, the *ref-alt* representation provided little improvement over *alt*. To control for the increased input vector size, we trained an equal-dimensionality control, *alt-alt-alt*, in which the alternate embedding was repeated three times so that the classifier input had the same input size as *ref-alt-diff*. Under the same LoRA configuration and training conditions, *alt-alt-alt* reached an AUROC of 0.848 and an AUPRC of 0.799, below both *alt* (0.860 and 0.811) and *ref-alt-diff* (0.868 and 0.834). These results suggest that explicitly incorporating the reference–alternate difference improves predictive performance beyond simply increasing the size of the input vector.

### 2.3 GLM-Missense improves substantially over zero-shot scoring

We combined the best-performing design choices from the preceding analyses to construct the final model. The final configuration used the NT-2 backbone, LoRA (*r* = 32, lr = 5×10*^−^*^5^), an MLP classifier head, and the variant-position *ref-alt-diff* input representation. We also expanded the training set to include variants classified in ClinVar as Pathogenic, Likely pathogenic, or Pathogenic/Likely pathogenic and as Benign, Likely benign, or Benign/Likely benign Landrum et al. (2018). We held out chromosomes 2, 3, 16, and 17 from training (Methods). We refer to this model as GLM-Missense. GLM-Missense achieved an AUROC of 0.896 and an AUPRC of 0.836 on a fixed subset of 5,000 variants from the validation set (Fig. S6).

Supervised fine-tuning improved over zero-shot scoring from the same NT-2 backbone. Zero-shot scores were computed using masked marginal log-likelihood, a standard language-model scoring strategy for mutation-effect prediction without task-specific supervision Meier et al. (2021). For this comparison, we used the ClinVar later-release test set, consisting of variants absent from the November 3, 2025 release but present in the September 23, 2026 release and located on the held-out chromo-somes (*n* = 5,780), thereby providing both temporal and chromosomal separation from model development (Methods). On this test set, GLM-Missense achieved an AUROC of 0.910 and an AUPRC of 0.867, compared with 0.751 AUROC and 0.700 AUPRC for zero-shot NT-2 scoring (Fig. S7). These results demonstrate that task-specific fine-tuning added substantial predictive signal beyond the general representations learned by the pretrained language model.

### 2.4 GLM-Missense contributes complementary predictive information beyond established predictors on held-out ClinVar variants

We next asked whether GLM-Missense provided novel information beyond established missense variant impact predictors. We used seven predictors calibrated by ClinGen for the PP3 and BP4 criteria Bergquist et al. (2025): BayesDel (without allele frequency) Feng (2017), MutPred2 Pejaver et al. (2020), REVEL Ioannidis et al. (2016), VEST4 Carter et al. (2013), AlphaMissense Cheng et al. (2023), ESM1b Brandes et al. (2023), and VARITY R Wu et al. (2021). These are the predictors currently recommended for computational evidence in clinical missense variant classification, and therefore the most clinically relevant baseline against which to ask whether GLM-Missense adds information. Unless otherwise noted, all comparisons were performed on the ClinVar later-release test variants with scores from every predictor (*n* = 3,327; 1,211 labeled as pathogenic and 2,116 labeled as benign). These variants are located on chromosomes that neither GLM-Missense nor the ensemble models had seen during training.

GLM-Missense showed the lowest concordance with the other predictors (Fig. 4A). Its pairwise Kendall’s *τ* correlation with the established predictors ranged from 0.43 to 0.54, whereas *τ* among the established predictors ranged from 0.53 to 0.80. Hierarchical clustering also placed GLM-Missense on a separate branch from the cluster containing the other predictors. This pattern suggested that GLM-Missense captured signal that was distinctive from the other predictors.

**Fig. 4.**
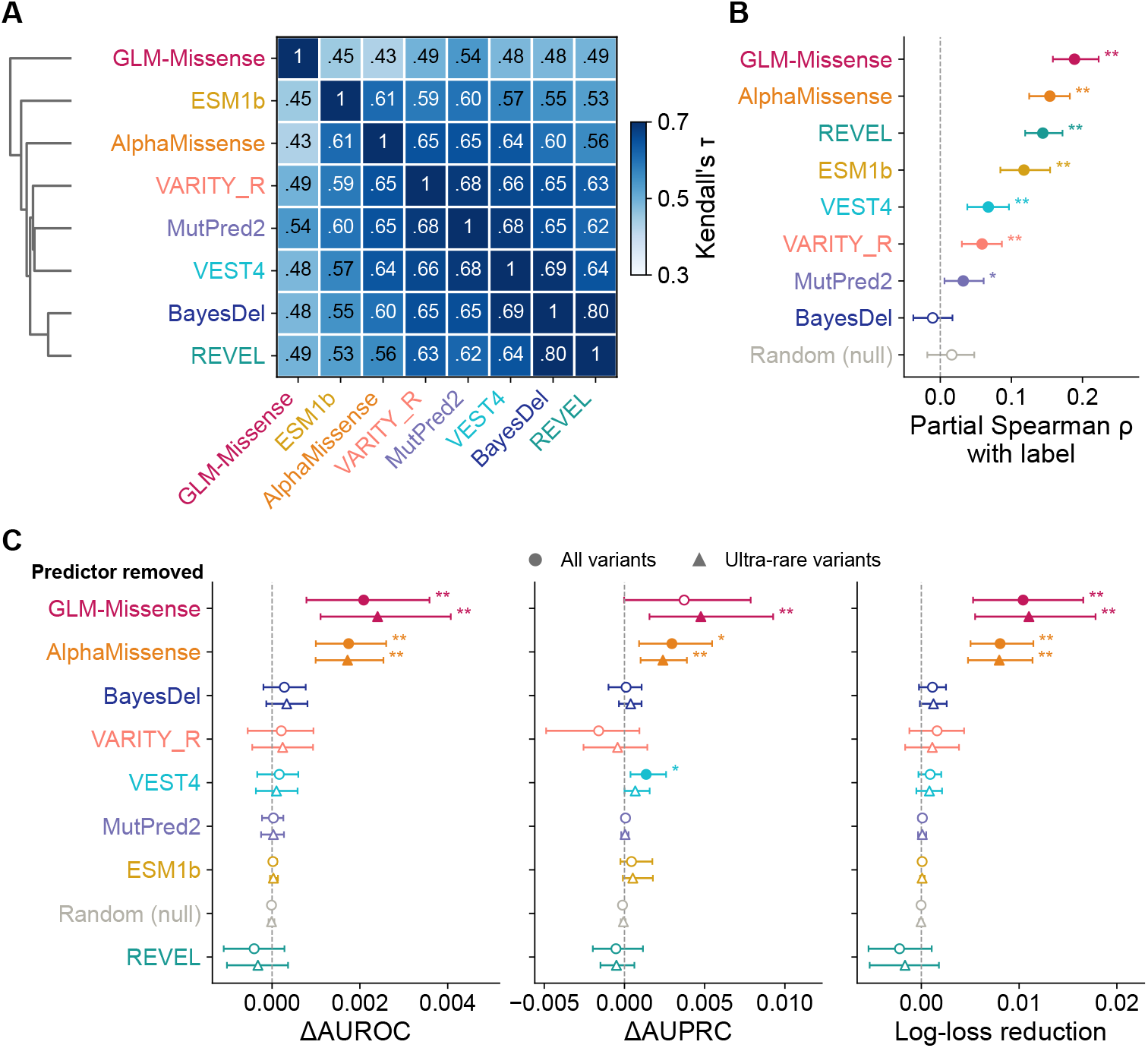
GLM-Missense provides complementary information beyond comparator predic-tors on held-out ClinVar variants. All panels use the ClinVar later-release test set with scores from all predictors (*n* = 3,327; 1,211 labeled as pathogenic and 2,116 labeled as benign), located on chromosomes 2, 3, 16, and 17, which were not used to train GLM-Missense or any ensemble. (**A**) Pair-wise Kendall’s *τ* between GLM-Missense and the seven comparator predictors, with average-linkage hierarchical clustering based on 1 *− τ* . GLM-Missense shows the lowest concordance with the other predictors. (**B**) Partial Spearman correlation between each predictor and the ClinVar label after con-trolling for the other seven predictor scores. Random (null), a score drawn from a uniform distribution between 0 and 1 for every variant, and its partial correlation is calculated after controlling for all eight predictors. GLM-Missense retains the strongest partial correlation. (**C**) Ensemble ablation analysis measuring the contribution of each predictor. The full ensemble, comprising the seven established predictors, GLM-Missense, and the random score, was compared with ensembles trained identically but excluding one predictor at a time. Positive values indicate that including the predictor improves the ensemble. Ensembles were trained on the later-release training set (*n* = 9,533) and evaluated on the full later-release test set (circles) and on ultra-rare variants in the later-release test set (triangles; gnomAD v4.1 joint allele frequency *<* 10^−4^ or absent from gnomAD; *n* = 3,082, 1,207 labeled as pathogenic and 1,875 labeled as benign). Error bars denote 95% CIs from 1,000 class-stratified boot-strap resamples, paired across rows. Filled symbols and stars denote Benjamini–Hochberg *q <* 0.05 within each panel (and, in C, within each variant stratum): ^∗^ *q <* 0.05, ^∗∗^ *q <* 0.01; open symbols are not statistically significant.

GLM-Missense’s distinctive signal was predictive rather than merely discordant, showing the strongest unique predictive signal among the evaluated methods. Following the rationale proposed by Lin et al. (unpublished manuscript), we quantified the unique predictive contribution of each method by asking how well each predictor remained associated with the ground-truth variant classification after accounting for the signal shared with the other predictor scores. Specifically, we computed the partial Spearman correlation between each predictor and the true label while controlling for the other seven predictor scores (Fig. 4B). The random score was drawn independently for each variant from a uniform distribution between 0 and 1, which by construction carries no information about pathogenicity. A larger positive value indicates that a method contributes useful information that is not already explained by the others. GLM-Missense showed the strongest partial correlation with pathogenicity (partial Spearman *ρ* = 0.189, 95% CI 0.158–0.223), followed by AlphaMissense (0.154), REVEL (0.144), and ESM1b (0.118). In contrast, BayesDel (−0.011; *q* = 0.42) and the random score (0.016; *q* = 0.42) showed no significant partial correlation. These results indicate that GLM-Missense contributes useful information not captured by established predictors, rather than uninformatively disagreeing with them.

Ensemble ablation analysis showed that GLM-Missense’s distinctive signal translated into practical predictive value. A predictor whose signal was already captured by the others would add little to an ensemble model, whereas a complementary predictor would still improve the ensemble performance. We trained an XGBoost ensemble Chen and Guestrin (2016), denoted MetaMissense, using scores from the seven established predictors and GLM-Missense. To avoid direct overlap between ensemble training and the variants used to develop GLM-Missense, we trained all ensembles on the ClinVar later-release training set rather than on the earlier-release set used for GLM-Missense model training (Methods). MetaMissense outperformed every individual predictor on the held-out test set, including GLM-Missense, which had lower standalone performance than the established predictors (Fig. S8). To quantify the contribution of each predictor, we compared the full ensemble with an otherwise identical ensemble trained without that predictor. We measured the differences in AUROC, AUPRC, and log-loss on the held-out ClinVar later-release test set using paired bootstrap 95% confidence intervals (CIs) and Benjamini–Hochberg-adjusted *q* values (Fig. 4C). Positive ΔAUROC, ΔAUPRC, and log-loss reduction indicate that including the predictor improves the ensemble. Including GLM-Missense increased AUROC by 0.0021 (95% CI 0.0008–0.0036; *q* = 0.0045) and reduced log-loss by 0.0104 (95% CI 0.0053–0.0166; *q* = 0.0045), both the largest improvements among all predictors. AlphaMissense was the only other predictor whose inclusion significantly improved both AUROC (0.0017) and log-loss (reduction 0.0081). Removing any of the remaining established predictors did not significantly change AUROC, suggesting that their information is largely shared with the other predictors. GLM-Missense also showed the largest improvement in AUPRC (ΔAUPRC 0.0037, 95% CI −0.00002 to 0.0079), although its improvement was not statistically significant (*q* = 0.16). The random score had no effect, as expected. Thus, including GLM-Missense improved both the ranking of variants and the predicted probabilities of an ensemble that already contained all seven comparator predictors.

In the ultra-rare variant setting, GLM-Missense remained the largest contributor to ensemble performance. Most variants in the ClinVar later-release test set are ultra-rare: 3,082 of 3,327 (93%) had a gnomAD v4.1 joint allele frequency below 10*^−^*^4^ or were absent from gnomAD. This included 1,207 of the 1,211 pathogenic variants and 1,875 of the 2,116 benign variants. On these ultra-rare variants, adding GLM-Missense increased AUROC by 0.0024 (95% CI 0.0011–0.0041) and reduced log-loss by 0.0110 (95% CI 0.0055–0.0178). Its contribution to AUPRC increased and became statistically significant (ΔAUPRC = 0.0048, 95% CI 0.0016–0.0093; *q* = 0.009; Fig. 4C, triangles). Restricting the analysis to ultra-rare variants excludes many higher-frequency benign variants that may be easier for some established predictors to identify, as methods such as AlphaMissense and VEST4 use population variants as benign training examples Cheng et al. (2023); Carter et al. (2013). Consistent with this possibility, the AUPRC contributions of AlphaMissense and VEST4 decreased on the ultra-rare variant sub-set, from 0.0030 to 0.0024 and from 0.0014 to 0.0007, respectively. The contribution of VEST4 was no longer statistically significant (*q* = 0.14). The remaining benign variants may therefore be more challenging to distinguish from pathogenic variants. Because AUPRC is sensitive to the ranking of benign variants among high-scoring predictions, the increased contribution of GLM-Missense suggests that it helps the ensemble assign lower pathogenicity scores to challenging ultra-rare benign variants, improving their separation from pathogenic variants.

We next asked whether the contribution of GLM-Missense could be explained by nucleotide-level conservation. As a standalone predictor, GLM-Missense outperformed the conservation scores phyloP 100-way vertebrate (AUROC 0.898, AUPRC 0.794), phyloP 470-way mammalian (0.884 and 0.757), and GERP++ (0.853 and 0.677) Pollard et al. (2010); Davydov et al. (2010) (Fig. S8). More importantly, when each score was added separately to an ensemble of the seven established predictors, GLM-Missense significantly improved its performance, whereas none of the conservation scores did (Fig. S9). One possible explanation is that nucleotide-level conservation is already incorporated into several established predictors, such as REVEL and Bayes-Del Ioannidis et al. (2016); Feng (2017). These results suggest that the complementary signal of GLM-Missense is not explained by the nucleotide-level conservation scores alone.

Finally, we asked whether the complementary contribution of GLM-Missense reflected its nucleotide-level input representation or its fine-tuning on recent ClinVar variants and classifications. Because GLM-Missense was fine-tuned on more recent ClinVar data, whereas established predictors were developed using earlier and different training datasets, GLM-Missense’s prediction could align more closely with recent ClinVar labels and diverge from established predictors, regardless of whether the model uses nucleotide- or protein-level inputs. To test this alternative explanation, we constructed a matched protein-model control, ESM1b-FT, by fine-tuning the protein language model ESM1b Rives et al. (2021) on the same training variants using the same downstream architecture, hyperparameters, and training procedure as GLM-Missense. On the same 5,000-variant validation subset, ESM1b-FT achieved an AUROC of 0.950 and an AUPRC of 0.923 (Fig. S10), outperforming GLM-Missense as a standalone predictor. Applying the same concordance and partial-correlation analyses to ESM1b-FT in place of GLM-Missense showed that ESM1b-FT was also less concordant with the established predictors and had the strongest partial correlation with pathogenicity labels (Fig. S11A, B). These results show that lower concordance and stronger partial correlation could also arise after fine-tuning a protein language model on recent ClinVar variants and classifications. However, the ensemble ablation revealed an important difference between the models: removing ESM1b-FT did not significantly change ensemble performance (Fig. S11C), whereas removing GLM-Missense did (Fig. 4C). The association of ESM1b-FT with the labels is therefore largely recoverable from the protein-centric predictors already in the ensemble, whereas GLM-Missense contributes information that other predictors in the ensemble cannot recover. Despite being trained on the same variants using the same settings, the two models made different contributions to ensemble performance, suggesting that training data alone cannot explain the contribution of GLM-Missense.

### 2.5 GLM-Missense retains its complementary predictive value on an independent functional assay dataset

ClinVar classifications partly depend on the computational evidence from the established predictors, which motivated us to investigate whether this may introduce circularity into our evaluation. Under the ACMG/AMP guidelines, submitters can assign the PP3 and BP4 criteria based on computational predictions, including those from the predictors evaluated here Richards et al. (2015); Pejaver et al. (2022); Bergquist et al. (2025). Consequently, evaluating these predictors against ClinVar classifications that were partly informed by their own scores may inflate their apparent performance, a form of circularity Grimm et al. (2015). ClinVar does not record in a structured form which ACMG/AMP criteria supported each classification, but the ClinGen Evidence Repository does so for expert-panel classifications Rehm et al. (2015). We therefore used the ClinGen Evidence Repository to identify later-release ClinVar variants with expert-panel classifications and scores available from all seven established predictors. We then recalculated each classification after removing PP3 and BP4 to identify variants whose classifications depend on computational evidence (Methods). Without these criteria, 71 of 176 (40%) pathogenic or likely pathogenic variants and 13 of 47 (28%) benign or likely benign variants would have been classified as variants of uncertain significance (VUS) (Fig. S12A, B). Excluding these 84 variants from the evaluation reduced the AUROC of all seven established predictors, significantly for BayesDel, REVEL, and VARITY R (*q <* 0.01; Fig. S12C). These results suggest that circularity may inflate the measured performance of the established pre-dictors on ClinVar variants. Because the contribution of GLM-Missense is measured relative to an ensemble of these predictors, whose performance is likely inflated, our estimates of its contribution on ClinVar variants are likely conservative.

To evaluate GLM-Missense on variants whose interpretation does not depend on computational predictions, we used a saturation genome editing (SGE) dataset for *RAD51D*, in which nearly all possible single-nucleotide variants were introduced into the endogenous locus in haploid HAP1 cells and classified by their effect on cell fit-ness Casadei et al. (2026). Because each variant is edited in its native genomic context and its transcripts are processed endogenously, this functional readout can reflect nucleotide-level effects on RNA processing in addition to the effect of the amino-acid substitution on the protein Findlay et al. (2018); Casadei et al. (2026). This dataset was particularly informative for two reasons. First, in addition to the functional read-out on cell fitness, Casadei et al. measured each variant’s frequency in RNA relative to its frequency in genomic DNA from the same cells. The resulting RNA score indicates whether transcripts carrying the variant are depleted. This RNA score provides experimental evidence, independent of computational predictions, of whether a variant’s effects may arise from altered RNA processing rather than solely from changes to the protein. Second, approximately 24% of loss-of-function *RAD51D* missense variants act through RNA-mediated mechanisms Casadei et al. (2026), so the dataset contains a substantial number of the variants for which we hypothesized that GLM-Missense could capture effects missed by protein-centric predictors. We used the functional classifications of 1,819 *RAD51D* missense variants: 227 were classified as functionally abnormal and 1,592 as functionally normal, excluding variants classified as indeterminate. *RAD51D* is located on chromosome 17, one of the held-out chromosomes, so none of these variants had been seen by GLM-Missense or the ensemble models during training. We evaluated ensembles trained on the ClinVar later-release training set on these variants, with ESM1b-FT replacing zero-shot ESM1b as one of the ensemble’s input predictors. Because ESM1b-FT was fine-tuned on the same variants and classifications as GLM-Missense using the same downstream settings, it provided a matched protein-level control for testing whether GLM-Missense’s complementary contribution could be explained by its fine-tuning data alone. It also served as a stronger standalone predictor than zero-shot ESM1b.

Consistent with the analysis based on ClinVar data, GLM-Missense retained its complementary contribution on the independent SGE dataset (Fig. 5). GLM-Missense again showed markedly lower concordance with the other predictors, with Kendall’s *τ* of 0.27–0.33, compared with 0.53–0.66 among the other predictors (Fig. 5A). It also showed the strongest partial correlation with the functional classifications (Fig. 5B). ESM1b-FT, in contrast, showed no unique association with the functional classifications. The ensemble ablation further demonstrated the contribution of GLM-Missense: the full ensemble reached an AUROC of 0.868, compared with 0.843 when GLM-Missense was removed (ΔAUROC = 0.0253, 95% CI 0.0166–0.0357; *q* = 0.0045). This was the largest change among all predictors; removing any other predictor changed AUROC by at most 0.0037. Including GLM-Missense also significantly reduced log-loss by 0.0333 (95% CI 0.0255–0.0413; *q* = 0.0011; Fig. 5C). However, the ensemble trained without ESM1b-FT performed better on the SGE dataset, with an AUROC increase of 0.0089 and a reduction in log-loss. Thus, although ESM1b-FT showed a strong partial correlation with ClinVar classifications, potentially reflecting its fine-tuning on ClinVar labels, this association did not persist on the independent SGE dataset. In contrast, GLM-Missense improved ensemble performance on both datasets, demonstrating that its complementary predictive value generalizes to independent experimental measurements of variant function.

**Fig. 5.**
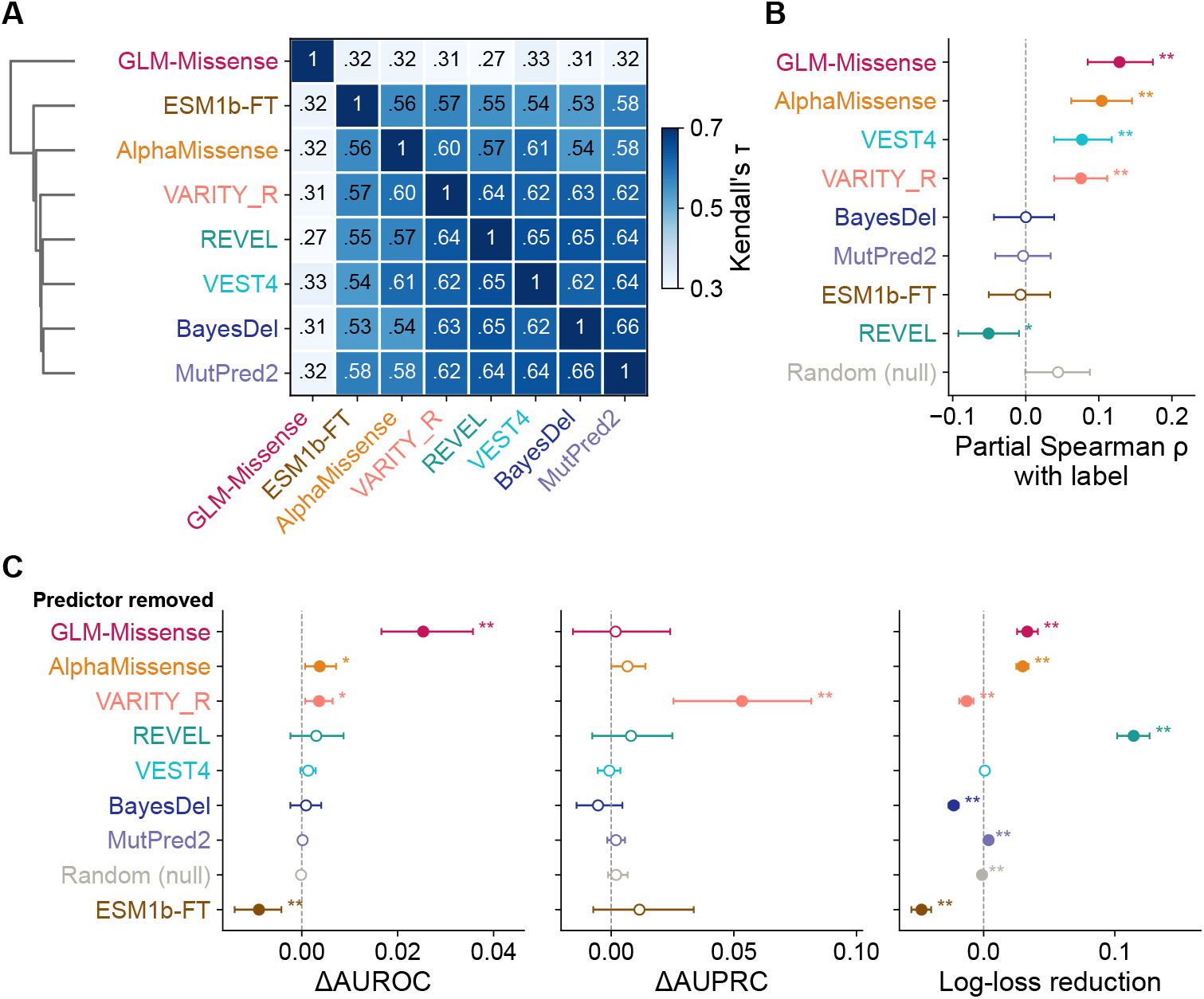
The contribution of GLM-Missense generalizes to independent *RAD51D* satu-ration genome editing data. All panels use the *RAD51D* missense variants classified by SGE as functionally abnormal (*n* = 227) or functionally normal (*n* = 1,592). Zero-shot ESM1b is replaced by ESM1b-FT among the established predictors. (**A**) Pairwise Kendall’s *τ* between GLM-Missense and the seven established predictors, with average-linkage hierarchical clustering based on 1 *− τ* . (**B**) Partial Spearman correlation between each predictor and the functional label after controlling for the other seven predictor scores; Random (null) is controlled for all eight predictors. (**C**) Ensemble ablation analysis, as in Fig. 4C. Ensembles were trained on the ClinVar later-release training set and evaluated on the *RAD51D*missense variants. Positive values indicate that including the predictor improves the ensemble. Error bars denote 95% CIs from 1,000 class-stratified bootstrap resamples, paired across rows. Filled symbols and stars denote Benjamini–Hochberg *q <* 0.05 within each panel: ^∗^ *q <* 0.05, ^∗∗^ *q <* 0.01; open symbols are not significant.

Ablation of several other predictors, including REVEL, BayesDel, and MutPred2, significantly changed ensemble log-loss but did not significantly affect AUROC. Unlike AUROC, which evaluates how well variants are ranked, log-loss depends on the pre-dicted probabilities and is sensitive to their calibration. The observed changes in log-loss may therefore reflect differences in how the ClinVar-trained ensembles’ pre-dicted probabilities align with the SGE functional classifications, rather than changes in their ability to distinguish between the two classes. The improvement in AUPRC from including GLM-Missense was not significant. The 95% CI for ΔAUPRC (−0.0156 to 0.0241) was approximately twice as wide as that for ΔAUROC, suggesting greater uncertainty in the estimated AUPRC change. This may partly reflect the limited number of functionally abnormal variants in the dataset (227; 12.5%). In addition, many functionally abnormal variants correctly classified by GLM-Missense were misclassified as functionally normal by most established predictors (see next section; Table 1).

**Table 1.** Functionally abnormal *RAD51D* missense variants resolved by GLM-Missense. The table includes all functionally abnormal *RAD51D* missense variants that were correctly classified by GLM-Missense but misclassified by at least five of the seven established predictors (with ESM1b-FT in place of ESM1b), together with functional and RNA scores, SpliceAI scores, exon numbers, exon-exon junction distances, and ESE hotspot status.

| Variant | Functional score | RNA score | SpliceAI score | Exon number | Junction dist | ESE hotspot |
|---|---|---|---|---|---|---|
| c.576G>T | −0.209 | −3.39 | 0.05 | 6 | 1 |  |
| c.576G>C | −0.229 | −8.08 | 0.08 | 6 | 1 |  |
| c.528C>G | −0.066 | −1.18 | 0.01 | 6 | 48 |  |
| c.528C>A | −0.069 | −1.13 | 0.01 | 6 | 48 |  |
| c.527T>G | −0.052 | −0.89 | 0.01 | 6 | 47 |  |
| c.480G>T | −0.222 | −2.46 | 0.9 | 5 | 1 |  |
| c.480G>C | −0.297 | −5.89 | 0.78 | 5 | 1 |  |
| c.422T>G | −0.164 | −2.81 | 0.04 | 5 | 59 |  |
| c.377A>C | −0.057 | 0.20 | 0.03 | 5 | 32 |  |
| c.341C>G | −0.071 | −2.01 | 0.25 | 4 | 5 |  |
| c.263G>T | −0.168 | −6.19 | 0.96 | 3 | 1 |  |
| c.263G>C | −0.289 | −7.53 | 0.96 | 3 | 1 |  |
| c.263G>A | −0.245 | −6.37 | 0.91 | 3 | 1 |  |
| c.262A>T | −0.313 | −7.54 | 0.76 | 3 | 2 |  |
| c.262A>G | −0.242 | −5.95 | 0.7 | 3 | 2 |  |
| c.262A>C | −0.254 | −7.75 | 0.56 | 3 | 2 |  |
| c.257T>G | −0.266 | −7.69 | 0.04 | 3 | 7 |  |
| c.251C>G | −0.214 | −4.55 | 0 | 3 | 13 |  |
| c.232T>G | −0.068 | −2.65 | 0 | 3 | 32 | Yes |
| c.229A>C | −0.056 | −1.97 | 0.03 | 3 | 35 | Yes |
| c.228G>T | −0.178 | −0.90 | 0.04 | 3 | 36 | Yes |
| c.226A>C | −0.151 | −5.33 | 0.03 | 3 | 38 | Yes |
| c.220G>C | −0.053 | −1.45 | 0.01 | 3 | 44 | Yes |
| c.220G>A | −0.105 | −1.25 | 0.01 | 3 | 44 | Yes |
| c.203G>C | −0.120 | −3.89 | 0 | 3 | 59 | Yes |
| c.202G>A | −0.187 | −2.47 | 0.42 | 3 | 58 | Yes |
| c.199A>T | −0.049 | −1.01 | 0.01 | 3 | 55 |  |
| c.185C>G | −0.055 | −1.08 | 0.05 | 3 | 41 |  |
| c.176C>G | −0.117 | −2.07 | 0.01 | 3 | 32 |  |
| c.167T>G | −0.071 | −2.49 | 0.04 | 3 | 23 |  |
| c.160A>G | −0.196 | −3.17 | 0.05 | 3 | 16 |  |
| c.146C>G | −0.058 | −3.59 | 0.03 | 3 | 2 |  |
| c.145G>T | −0.142 | −2.10 | 0.53 | 3 | 1 |  |
| c.144G>T | −0.126 | −1.87 | 0.63 | 2 | 1 |  |
| c.144G>C | −0.109 | −4.63 | 0.64 | 2 | 1 |  |
| c.82G>T | −0.174 | 2.32 | 0.93 | 1 | 1 |  |
| c.82G>C | −0.145 | −5.58 | 0.95 | 1 | 1 |  |
| c.82G>A | −0.144 | −3.78 | 0.88 | 1 | 1 |  |

**Variant,** HGVS coding variant relative to ENST00000345365.11; **Functional score,** SGE-derived measure of the variant's effect on cell fitness Casadei et al. (2026); **RNA score,** log<sub>2</sub> ratio of the variant's frequency in RNA to its frequency in genomic DNA Casadei et al. (2026); **SpliceAI score,** maximum SpliceAI delta score across acceptor gain, acceptor loss, donor gain, and donor loss; **Exon number,** exon in which the variant lies; **Junction dist,** distance (nt) of the variant to the nearest exon-exon junction; **ESE hotspot,** Yes if the variant is located in the exonic splicing enhancer (ESE)-rich interval c.202\_235, whose deletion completely abolished the full-length transcript in the minigene assays of Bueno-Martínez et al. (2021). All exon 3 variants except those at its first two or last three nucleotides lie within one of four overlapping intervals (c.147\_176, c.172\_206, c.202\_235 and c.231\_260) whose deletion also impaired exon 3 recognition in the same assays.

Ranking these functionally abnormal variants higher could improve AUROC without a comparable improvement in AUPRC, which is particularly sensitive to precision among the highest-scoring variants. In contrast, including VARITY R significantly improved AUPRC on the SGE dataset but not on ClinVar. VARITY was the only comparator method trained in part on multiplexed functional assay data Wu et al. (2021), potentially explaining its stronger contribution to AUPRC on the SGE dataset.

Together, these results show that GLM-Missense captures information overlooked by established predictors that generalizes from clinical classifications to independently measured variant effects. This information was not captured by ESM1b-FT, a protein language model fine-tuned on the same variants as GLM-Missense. Because the two models share the same training data but differ only in their input representation, the additional information captured by GLM-Missense is likely attributable to its nucleotide-level representation rather than to its training data.

### 2.6 Splicing disruption accounts for part of the complementary signal of GLM-Missense

To identify what nucleotide-level information GLM-Missense captures, we first examined variants in the ClinVar later-release test set that were correctly classified by GLM-Missense but misclassified by at least five of the seven established predictors, using the ClinGen-calibrated PP3 and BP4 Supporting thresholds of each predictor (Methods). We refer to these as GLM-Missense-resolved variants. Because missense-annotated variants can affect function through splicing disruption in addition to, or instead of, the amino-acid substitution, we compared the predicted splicing impact of resolved and other variants using the maximum SpliceAI delta score across acceptor gain, acceptor loss, donor gain, and donor loss Jaganathan et al. (2019). The 31 pathogenic or likely pathogenic variants resolved by GLM-Missense had significantly higher SpliceAI scores than the remaining pathogenic or likely pathogenic variants (*p* = 0.0045; Fig. S13A). Of these 31 variants, 14 were located within two nucleotides of an exon boundary. Thirteen of these 14 had a maximum SpliceAI delta score of at least 0.2, including all six variants with splicing-related evidence in their Clin-Var records (Table S2). In contrast, no significant difference in SpliceAI scores was observed for the 41 pathogenic or likely pathogenic variants resolved by ESM1b-FT (*p* = 0.12; Fig. S13B), or for the benign or likely benign variants resolved by either model. The enrichment for predicted splicing disruption is therefore specific to GLM-Missense rather than a general property of models fine-tuned on the same variants as GLM-Missense.

Because ClinVar records provide experimental evidence on the mechanism for only a few of the GLM-Missense-resolved ClinVar variants, we next examined GLM-Missense-resolved variants in the *RAD51D* SGE data. In addition to the functional classifications, this dataset provides an RNA score readout, defined as the log_2_ ratio of each variant’s frequency in RNA to its frequency in genomic DNA from the same cells Casadei et al. (2026). Negative RNA scores indicate reduced abundance of transcripts carrying the variant. The RNA score therefore serves as an experimental proxy for splicing-related and other RNA-mediated effects. Among the 227 functionally abnormal variants, the 38 functionally abnormal variants resolved by GLM-Missense had significantly lower RNA scores than the remaining 189, with median scores of −2.73 and −0.35, respectively (*p* = 1.6 × 10*^−^*^8^; Fig. 6A). No significant difference was observed between GLM-Missense-resolved and other functionally normal variants (*p* = 0.51). Together, these findings suggest that GLM-Missense captures information related to RNA-mediated effects, as functionally abnormal variants resolved by the model had significantly reduced RNA abundance.

**Fig. 6.**
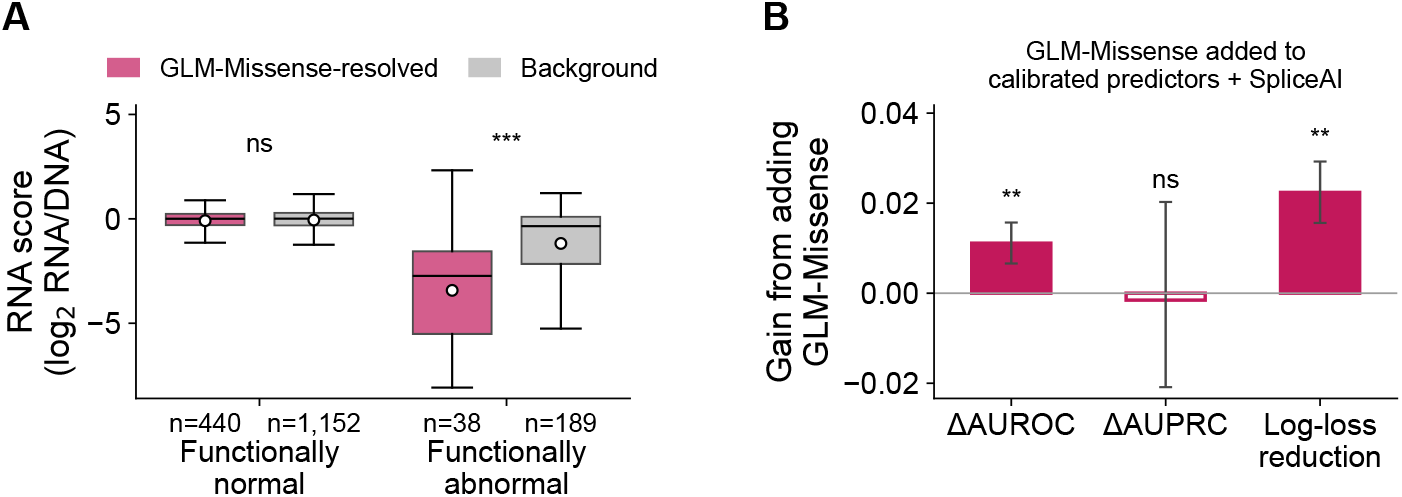
GLM-Missense captures RNA-mediated effects on *RAD51D* SGE data, includ-ing information missed by SpliceAI. **(A)** RNA scores (log_2_ ratio of the variant frequency in RNA to that in genomic DNA) of GLM-Missense-resolved and background *RAD51D* variants, shown separately for functionally normal (resolved *n* = 440, background *n* = 1,152) and functionally abnor-mal (resolved *n* = 38, background *n* = 189) variants. GLM-Missense-resolved variants were correctly classified by GLM-Missense but misclassified by at least five of the seven established predictors (with ESM1b-FT in place of ESM1b); background variants comprise all remaining variants within each functional class. Boxes represent the interquartile range with lines indicating medians, whiskers extending to the most extreme values within 1.5 times the interquartile range, and white circles indi-cating means. Values beyond the whiskers are not shown. Two-sided Mann–Whitney *U*test: ns, not significant; ^∗∗∗^ *p <* 0.001. **(B)** Gain from adding GLM-Missense to an ensemble of the seven established predictors (with ESM1b-FT) and SpliceAI. Ensembles with and without GLM-Missense were trained identically on the ClinVar later-release training set and evaluated on the *RAD51D* SGE vari-ants (*n* = 1,819). Bars show ΔAUROC, ΔAUPRC, and log-loss reduction; error bars denote 95% CIs from 1,000 class-stratified paired bootstrap resamples. Two-sided bootstrap *p*: ns, not significant; ^∗∗^ *p <* 0.01.

We next asked whether GLM-Missense still adds information to an ensemble that already includes a well-established splicing predictor. We compared two ensembles trained identically on ClinVar data, both containing the seven established predictors (with ESM1b-FT replacing ESM1b) and SpliceAI, but only one including GLM-Missense. On the *RAD51D* SGE dataset, adding GLM-Missense increased AUROC from 0.867 to 0.878 (ΔAUROC = 0.0111; 95% CI 0.0066–0.0157; *p* = 0.001), and significantly reduced the log-loss by 0.0224 (95% CI 0.0156–0.0292; *p* = 0.001), whereas the change in AUPRC was not significant (Fig. 6B). Thus, the contribution of GLM-Missense is not fully explained by the splicing effects captured by SpliceAI.

To investigate the potential sources of GLM-Missense’s complementary predictive information beyond SpliceAI, we examined the functionally abnormal variants resolved by GLM-Missense (Table 1). Notably, 23 of the 38 functionally abnormal variants resolved by GLM-Missense were located in exon 3 of *RAD51D* (Table 1). Previous studies indicate that this exon is particularly susceptible to splicing disruption. Casadei et al. found that exons 3 and 4 contain the highest concentration of RNA-mediated loss-of-function variants in *RAD51D* Casadei et al. (2026). Minigene assays by Bueno-Martínez et al. further showed that correct inclusion of exon 3 in the mature transcript depends not only on its canonical splice sites but also on exonic splicing enhancers (ESEs) within the exon Bueno-Martínez et al. (2021). ESEs are short sequence motifs that recruit splicing activators, primarily serine/arginine-rich (SR) proteins, and help the splicing machinery recognize the adjacent splice sites Cartegni et al. (2002); Fair-brother et al. (2002). Deletion of any of four overlapping intervals spanning exon 3 (c.147 176, c.172 206, c.202 235, and c.231 260) from the minigene impaired inclusion of the exon, indicating that ESEs are present throughout the exon Bueno-Martínez et al. (2021). Specifically, deleting the ESE-rich interval c.202 235 abolished the full-length transcript completely. Exon 3 skipping shifts the reading frame and introduces a premature termination codon, potentially leading to nonsense-mediated mRNA decay Maquat et al. (1981) and loss of *RAD51D* function. This mechanism is consistent with the reduced RNA scores of these variants. Missense-annotated variants in exon 3 may therefore cause loss of *RAD51D* function and reduced RNA abundance by disrupting exonic splicing regulatory elements, rather than through the effects of the amino-acid substitution.

Importantly, ESEs can be disrupted by variants located away from exon boundaries. Such variants are difficult for protein-level predictors to detect, as these predictors focus mainly on the effect of the amino-acid substitution, and they can also be missed by splicing predictors. Of the 23 GLM-Missense-resolved variants in exon 3, 14 variants are located more than 10 nucleotides from the nearest exon boundary. Of these 14 variants, eight fell within the ESE-rich interval c.202 235, while the remaining six fell within the other intervals whose deletion also impaired exon 3 inclusion. All 14 variants had negative RNA scores, indicating reduced transcript abundance and sup-porting the possibility that these variants disrupt exonic splicing regulatory elements. Notably, SpliceAI missed almost all of them: 13 of the 14 variants have maximum SpliceAI delta scores below 0.2, meaning that SpliceAI predicts them to be unlikely to alter splicing. These findings suggest that the splicing-related information captured by GLM-Missense is not limited to variants near splice junctions but may also include disruption of exonic splicing regulatory elements deeper within exons. Such effects may be missed even by a state-of-the-art splicing predictor, potentially explaining part of GLM-Missense’s complementary contribution beyond SpliceAI.

### 2.7 Codon-level effects on mRNA stability may also contribute to the complementary signal of GLM-Missense

Because GLM-Missense evaluates variants in their nucleotide sequence context, it may capture nucleotide-level effects beyond splicing disruption. However, such effects may be subtle and difficult to detect among the relatively small number of labeled variants resolved by GLM-Missense. We therefore examined the much larger set of ClinVar mis-sense variants of uncertain significance (VUS) on the held-out chromosomes. Although VUS lack definitive classifications, comparing variants on which GLM-Missense dis-agrees with established predictors against matched controls may reveal nucleotide-level features associated with its complementary signal.

We identified discordant VUS for which GLM-Missense made a confident pathogenic (score ≥ 0.8) or benign (score ≤ 0.2) prediction opposite to the calls of at least five of the seven established predictors. For each discordant VUS, we selected a matched control using three criteria: the control variant (1) was from the same gene, (2) received the same benign or pathogenic prediction from the same number of established predictors, but an opposite confident prediction from GLM-Missense, and (3) was the nearest eligible VUS in genomic position.

We compared discordant VUS with their matched controls using three nucleotide-level annotations, beginning with the maximum SpliceAI delta score as a positive control because the preceding analyses of ClinVar and SGE variants had identified an association between GLM-Missense-resolved variants and splicing-related effects. Consistent with that finding, VUS that GLM-Missense called pathogenic against most established predictors (*n* = 1,254 pairs) had significantly higher SpliceAI scores than their matched controls (*q* = 1.8×10*^−^*^18^; Fig. 7A), whereas VUS it called benign against most established predictors (*n* = 2,675 pairs) had lower SpliceAI scores (*q* = 0.0055).

**Fig. 7.**
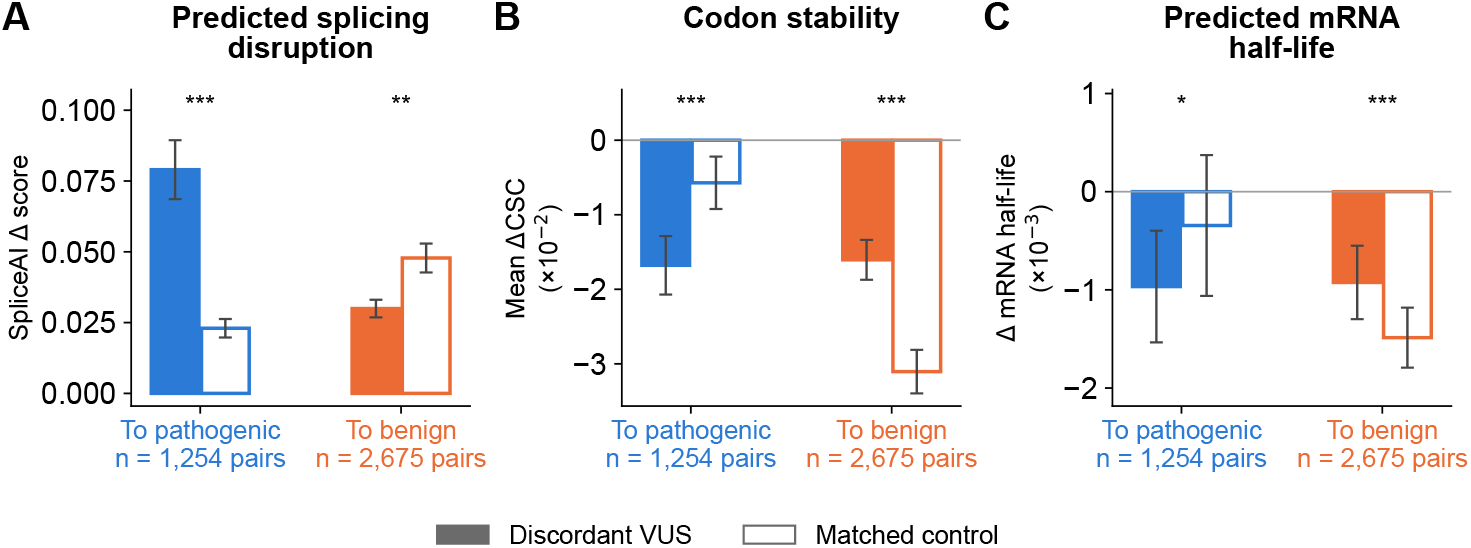
VUS on which GLM-Missense disagrees with most established predictors differ from matched controls in splicing and mRNA-stability annotations. Discordant VUS (filled bars) were missense VUS on the held-out chromosomes for which GLM-Missense made a confident pathogenic (score *≥* 0.8) or benign (score *≤* 0.2) prediction opposite to at least five of the seven established predictors. Each discordant VUS was paired with a matched control (open bars) from the same gene that received the same benign or pathogenic prediction from the same number of estab-lished predictors, but a confident prediction from GLM-Missense in agreement with the established predictors. The nearest eligible VUS in genomic position was selected as the control. *To pathogenic* denotes discordant VUS that GLM-Missense predicted to be pathogenic but at least five established predictors predicted to be benign (*n* = 1,254 pairs); *To benign* denotes the reverse (*n* = 2,675 pairs). **(A)** Maximum SpliceAI delta score. **(B)** Mean change in codon stability coefficient (ΔCSC, alternate minus reference codon), averaged over six human CSC datasets Wu et al. (2019). Negative values indicate a change toward a less optimal codon. **(C)** Change in mRNA half-life predicted by Saluki (alternate minus reference transcript). Bars represent means, and error bars indicate 95% confidence intervals of the means. Two-sided Wilcoxon signed-rank tests on the pairs, with Benjamini–Hochberg adjustment over the six tests shown: ^∗^ *q <* 0.05, ^∗∗^ *q <* 0.01, ^∗∗∗^ *q <* 0.001.

We next examined two annotations related to mRNA stability. Single-nucleotide substitutions can alter codon optimality, which influences mRNA stability and translation and may thereby affect gene expression Presnyak et al. (2015); Wu et al. (2019). Because codon optimality depends on the nucleotide sequence rather than solely on the encoded amino acid, this information is not directly available to protein-sequence-based models. We assessed potential effects on mRNA stability using the change in codon stability coefficient (ΔCSC) and the change in predicted mRNA half-life. The CSC of a codon measures the correlation, across mRNAs, between its frequency and mRNA stability Presnyak et al. (2015). We averaged ΔCSC (alternate minus reference codon) across six human CSC datasets Wu et al. (2019), with negative values indicating a change toward a less optimal codon. The change in mRNA half-life between the alternate and reference transcripts was predicted by Saluki, a model that pre-dicts mRNA half-life from mRNA sequence Agarwal and Kelley (2022). Compared with their matched controls, VUS that were called pathogenic by GLM-Missense had more negative ΔCSC values (*q* = 4.3 × 10*^−^*^5^; Fig. 7B) and more negative predicted changes in mRNA half-life (*q* = 0.019; Fig. 7C). Conversely, VUS called benign by GLM-Missense had less negative ΔCSC values (*q* = 5.7 × 10*^−^*^14^) and less negative predicted changes in mRNA half-life (*q* = 6.6 × 10*^−^*^5^) than their controls. We also applied the same analysis to ESM1b-FT, a protein language model fine-tuned on the same variants as GLM-Missense. In contrast, none of the three annotations differed significantly between ESM1b-FT-discordant VUS and their matched controls in either prediction direction (Fig. S14).

The associations observed for GLM-Missense but not for ESM1b-FT, despite fine-tuning on the same variants, together with the same gene matching control, suggest that these associations are unlikely to be explained by the characteristics of training data or by gene identity, but may instead reflect nucleotide-level properties beyond those captured by protein-based models. Although VUS lack definitive classifications and these results therefore cannot establish whether individual predictions are correct, they suggest that, beyond splicing disruption, codon-level effects on mRNA stability may also contribute to the complementary predictive information captured by GLM-Missense.

## 3 Discussion

### 3.1 Multi-species pretraining and variant-centered representations capture core biological properties of missense variants

Multi-species pretraining was a major factor affecting performance in the frozen-backbone setting (Fig. 2). NT-2 was pretrained on genomes from 850 species spanning diverse phyla Dalla-Torre et al. (2025), whereas NT-1 was pretrained only on human genomes. The advantage of NT-2 is consistent with the central role of evolutionary constraint in missense variant interpretation Cooper and Shendure (2011); Capriotti and Fariselli (2022): positions conserved across species are often functionally constrained, so variants at those positions are more likely to be deleterious, and such patterns can-not be observed in human sequences alone. NT-1 and NT-2 also differ in architecture and pretraining hyperparameters, making it difficult to isolate the effect of pretraining corpus diversity. However, an independent comparison reported in Genomic Touch-stone found that a multi-species Nucleotide Transformer model generally outperformed a human-only counterpart with the same architecture and model size Wang et al. (2025). Together, these findings suggest that pretraining corpus diversity may be a key driver of genomic language model performance in missense variant impact prediction. Variant-position embeddings were consistently more informative than pooled sequence representations (Fig. 2). This pattern is expected for missense variants, whose pathogenic effects often depend on the altered residue and its immediate molecular context, including effects on stability, catalytic function or molecular interfaces Gerasi-mavicius et al. (2020, 2022). In NT-2, the representation at the variant position already integrates surrounding sequence context through the self-attention mechanism dur-ing pretraining Dalla-Torre et al. (2025). In contrast, mean pooling over thousands of tokens, many of which correspond to positions unrelated to the affected codon (e.g., intronic or intergenic regions), would likely dilute this variant-specific information.

### 3.2 LoRA preserves pretrained representations while enabling effective task adaptation

LoRA was effective for adapting NT-2 to missense variant impact prediction. It achieved the highest validation AUROC and AUPRC, whereas full fine-tuning showed weaker generalization consistent with overfitting (Fig. S4). This contrast is notable because the fine-tuning dataset was small relative to the model size. Full fine-tuning grants the optimizer enough degrees of freedom to fit training-set-specific patterns that may not generalize. Compounding this, NT-2 has already captured essential genomic features during pretraining on billions of nucleotides across 850 species Dalla-Torre et al. (2025), such as evolutionary constraints that are directly relevant to variant pathogenicity. Updating all parameters therefore risks overwriting these broadly use-ful representations in favor of spurious patterns in the comparatively small labeled set. Such loss of pretrained knowledge is broadly termed catastrophic forgetting McCloskey and Cohen (1989); Kirkpatrick et al. (2017).

LoRA reduces the risk of overfitting by restricting how the pretrained weights can be updated during fine-tuning. It freezes the pretrained weights *W*_0_ and learns only low-rank perturbations, Δ*W* = *BA*, with rank *r* = 32 ≪ *d* = 1,024. This low-rank constraint limits the model’s capacity to memorize training-specific patterns while still permitting meaningful task adaptation, consistent with the “learns less, forgets less” characterization of LoRA in large language models Biderman et al. (2024). Importantly, in our experiments, LoRA not only avoided overfitting but also achieved the highest validation performance among all runs, surpassing every full fine-tuning configuration in both AUROC and AUPRC. LoRA is often framed as a tradeoff in which preserving base-model knowledge comes at some cost to peak task performance. Our result instead suggests that when pretrained representations are relatively well aligned with the downstream task, low-rank adaptation acts as a useful inductive bias rather than a limitation.

LoRA experiments also showed an advantage in sample efficiency. LoRA achieved 0.860 validation AUROC using only 5k training samples, within 0.011 AUROC of the frozen-backbone baseline trained on the 44k samples (0.871 AUROC), while updating only about 0.8% of the backbone parameters. This efficiency is especially relevant for clinical genetics, where labeled data are often scarce. Rare diseases involve small patient cohorts, understudied populations are affected by ascertainment bias, and newly implicated genes often have few annotated variants Landrum et al. (2018). Our results therefore suggest that LoRA can leverage and adapt pretrained genomic representations efficiently even with limited data.

Fine-tuning was also substantially more effective than zero-shot scoring from the same NT-2 backbone (Fig. S7). This difference is expected because masked language-model likelihood measures how well an alternate nucleotide fits the pretrained sequence distribution, rather than whether the resulting variant is clinically pathogenic or benign Meier et al. (2021). In long genomic windows, this likelihood can reflect many sequence constraints, including local nucleotide composition, repetitive sequence, and other contextual patterns that are not specific to the functional consequence of a single missense substitution. Fine-tuning therefore serves an essential role: it redirects the general sequence representation learned during pretraining toward the clinical classification boundary of interest.

### 3.3 GLM-Missense captures distinctive nucleotide-level information overlooked by established missense variant impact predictors

Most established missense variant impact predictors, including the seven comparator predictors in this study, focus primarily on the protein-level consequences of amino-acid substitutions, using protein sequence, structure, and evolutionary conservation either directly or through the scores of other predictors Brandes et al. (2023); Cheng et al. (2023); Carter et al. (2013); Pejaver et al. (2020); Ioannidis et al. (2016); Feng (2017); Wu et al. (2021). Since most pathogenic missense variants are thought to cause disease through the effect of the amino-acid substitution on protein structure or function, protein-centric predictors likely capture much of the information relevant to pathogenicity. GLM-Missense, in contrast, has to infer protein-level consequences indirectly from DNA sequence, so it is not surprising that it underperformed these predictors as a standalone classifier (Fig. S8).

However, standalone performance does not necessarily reflect the value a predictor adds to an ensemble. Removing GLM-Missense from an ensemble that already contained seven established predictors reduced performance more than removing any of them, both on held-out ClinVar variants (Fig. 4C) and on the *RAD51D* SGE data (Fig. 5C). In contrast, most of the established predictors could each be removed with-out a significant reduction in ensemble performance, indicating that much of their information is shared, likely because they draw on similar protein-level features. Thus, a predictor with lower standalone performance can still provide complementary infor-mation that improves predictions beyond what stronger individual predictors achieve together. This suggests that some of the remaining errors in current missense variant impact prediction may reflect biological information not adequately captured by exist-ing methods, rather than only irreducible uncertainty. Furthermore, the contribution of GLM-Missense may be underestimated on the ClinVar dataset. ClinVar classifications can incorporate computational evidence from the established predictors, which may favor these predictors in evaluations based on ClinVar (Fig. S12). The complementary contribution of GLM-Missense on the independent SGE dataset, whose functional classifications do not rely on computational predictions, provides further evidence that its predictive value generalizes beyond ClinVar data.

The most plausible reason GLM-Missense adds distinctive information is its nucleotide-level representation. ESM1b-FT, a protein language model fine-tuned on the same training variants as GLM-Missense, did not reproduce its contribution (Figs. 5C and S11C). As discussed in the Background, missense-annotated variants can also act through their nucleotide sequence context, for example, by altering splicing or codon usage. Such effects are not directly represented by models that operate solely on protein sequences. GLM-Missense instead operates directly on DNA sequence and can, in principle, represent both protein-level consequences and nucleotide-level effects. Our analyses identified splicing disruption as one such effect and suggested codon-level effects on mRNA stability as another.

Splicing disruption by missense-annotated variants is well recognized, particularly at positions adjacent to canonical splice sites Soemedi et al. (2017); Jaganathan et al. (2019), and many ClinVar variants resolved by GLM-Missense were located near exon boundaries (Table S2). The *RAD51D* analysis suggests that GLM-Missense may also capture splicing-related effects deeper within exons, away from canonical splice sites. Most functionally abnormal *RAD51D* variants resolved by GLM-Missense were located in exon 3, whose inclusion depends on exonic splicing enhancers present throughout the exon Bueno-Martínez et al. (2021). Many of these variants were far from the exon boundaries, yet transcripts carrying them were substantially less abundant (Fig. 6A; Table 1). Such variants fall into a potential blind spot in current computational approaches to missense variant impact prediction. The established pre-dictors, which focus on the effect of the amino-acid substitution, are not designed to capture such effects on splicing. SpliceAI also missed nearly all of these variants: their scores were below 0.2, the threshold recommended by the ClinGen Sequence Vari-ant Interpretation (SVI) Splicing Subgroup for using SpliceAI as evidence of splicing impact. Our findings are consistent with those of the ClinGen SVI Splicing Subgroup, which reported that SpliceAI performed less well on exonic variants, potentially due to its limited ability to predict variants disrupting exonic splicing enhancers Walker et al. (2023). The splicing signal captured by GLM-Missense therefore extends beyond splice-site-adjacent positions to exonic splicing regulatory elements, which may be overlooked by both protein-centric predictors and a state-of-the-art splicing predictor. Codon-level effects on mRNA stability represent another potential source of complementary information, although the supporting evidence is less direct. Protein-centric models assign the same score to every nucleotide change that produces the same amino-acid substitution, even though these changes may introduce codons that differ in optimality. Codon optimality influences mRNA stability and translation in human cells Presnyak et al. (2015); Wu et al. (2019). Moreover, nucleotide changes that alter codon usage without changing the encoded protein have been implicated in human disease Sauna and Kimchi-Sarfaty (2011). In our VUS analysis, disagree-ment between GLM-Missense and the established predictors was associated with codon optimality and predicted mRNA half-life. VUS that GLM-Missense predicted to be pathogenic, against most established predictors, introduced less optimal codons and were predicted to shorten mRNA half-life, whereas those it predicted to be benign showed the opposite pattern (Fig. 7B, C). These associations were not observed for ESM1b-FT, a protein-model control matched to GLM-Missense that does not capture the codon-level information (Fig. S14). A change toward a less optimal codon could reduce mRNA stability and thereby protein abundance, potentially affecting function in dosage-sensitive genes. However, the association with GLM-Missense predictions does not establish whether changes in mRNA stability contribute to pathogenicity, and the observed differences between discordant VUS and their controls were relatively small (Fig. 7B, C). Whether individual codon changes alter mRNA stability sufficiently to contribute to disease therefore remains to be established. Together, these findings show that a fine-tuned genomic language model can capture nucleotide-level information overlooked by current missense variant impact predictors.

## 4 Conclusion

We systematically explored how a pretrained genomic language model can be adapted for clinical missense variant impact prediction. Preserving the representation at the variant position, pretraining on multi-species genomes, explicitly contrasting the reference and alternate alleles, and adapting the model backbone with LoRA yielded the best performance. The resulting model, GLM-Missense, substantially outperformed zero-shot scoring from the same pretrained backbone.

GLM-Missense captured nucleotide-level information overlooked by established missense variant impact predictors. MetaMissense, an ensemble that combines GLM-Missense with the seven comparator predictors, outperformed each individual pre-dictor. Removing GLM-Missense from the ensemble led to the greatest decrease in performance of any predictor. This complementary contribution of GLM-Missense was observed on both held-out ClinVar variants and an independent *RAD51D* saturation genome editing dataset, whose functional classifications do not depend on computational predictions. A protein language model fine-tuned on the same training variants as GLM-Missense did not reproduce this contribution. The nucleotide-level information captured by GLM-Missense appears to reflect, in part, effects on splicing. This includes disruption of exonic splicing regulatory elements located away from canonical splice sites, which can be overlooked by protein-centric predictors and a state-of-the-art splicing predictor. Analysis of variants of uncertain significance further suggested that GLM-Missense may also capture codon-level effects on mRNA stability.

These findings have implications for clinical variant interpretation, where computational evidence under the ACMG/AMP PP3 and BP4 criteria relies heavily on protein-centric predictors. They suggest that some of the residual errors of established protein-centric predictors may reflect nucleotide-level information not adequately captured by existing methods, and that fine-tuned genomic language models can recover part of this information. Our results also highlight the importance of evaluating newly developed predictors not only by their standalone performance but also by their contributions to existing ensembles. By capturing information beyond the consequences of amino-acid substitutions, genomic language models may help improve the interpretation of missense-annotated variants through a broader consideration of their potential functional effects.

## 5 Methods

### 5.1 ClinVar sets, label mappings, and chromosome-exclusive splits

We curated GRCh38 missense variants from ClinVar Landrum et al. (2018, 2020) and used two binary label mappings. Under the *strict* mapping, we retained only variants classified as *Benign* or *Pathogenic*. Under the *expanded* mapping, the benign class comprised *Benign*, *Likely benign*, and *Benign/Likely benign*, and the pathogenic class comprised *Pathogenic*, *Likely pathogenic*, and *Pathogenic/Likely pathogenic*. Both mappings excluded *Conflicting classifications*, *Uncertain significance*, and all other ClinVar categories. The strict mapping has the clearest labels and was used for architecture exploration. The expanded mapping was used to train GLM-Missense and ESM1b-FT and for all downstream analyses.

We defined two non-overlapping sets by ClinVar release. The *earlier-release set* contained variants with a benign or pathogenic classification in the November 3, 2025 release. The *later-release set* contained variants with such a classification in the September 23, 2026 release that were absent from all six benign and pathogenic classes (B, LB, B/LB, P, LP, P/LP) of the earlier release. Variants were matched between releases by chromosome, position, reference allele, and alternate allele, not by ClinVar variation or submission identifiers. A resubmitted variant therefore could not appear in both sets. Every later-release variant was first reported in ClinVar after Novem-ber 3, 2025, after the publication of all established predictors evaluated here. This temporal separation prevented overlap between model development and evaluation.

All training, validation, and test splits were chromosome-exclusive. This design pre-vents variants from the same gene, or from nearby correlated positions, from appearing in both training and evaluation sets. Table 2 summarizes all sets, their chromosomes, and their numbers of variants in each ClinVar class.

**Table 2.**
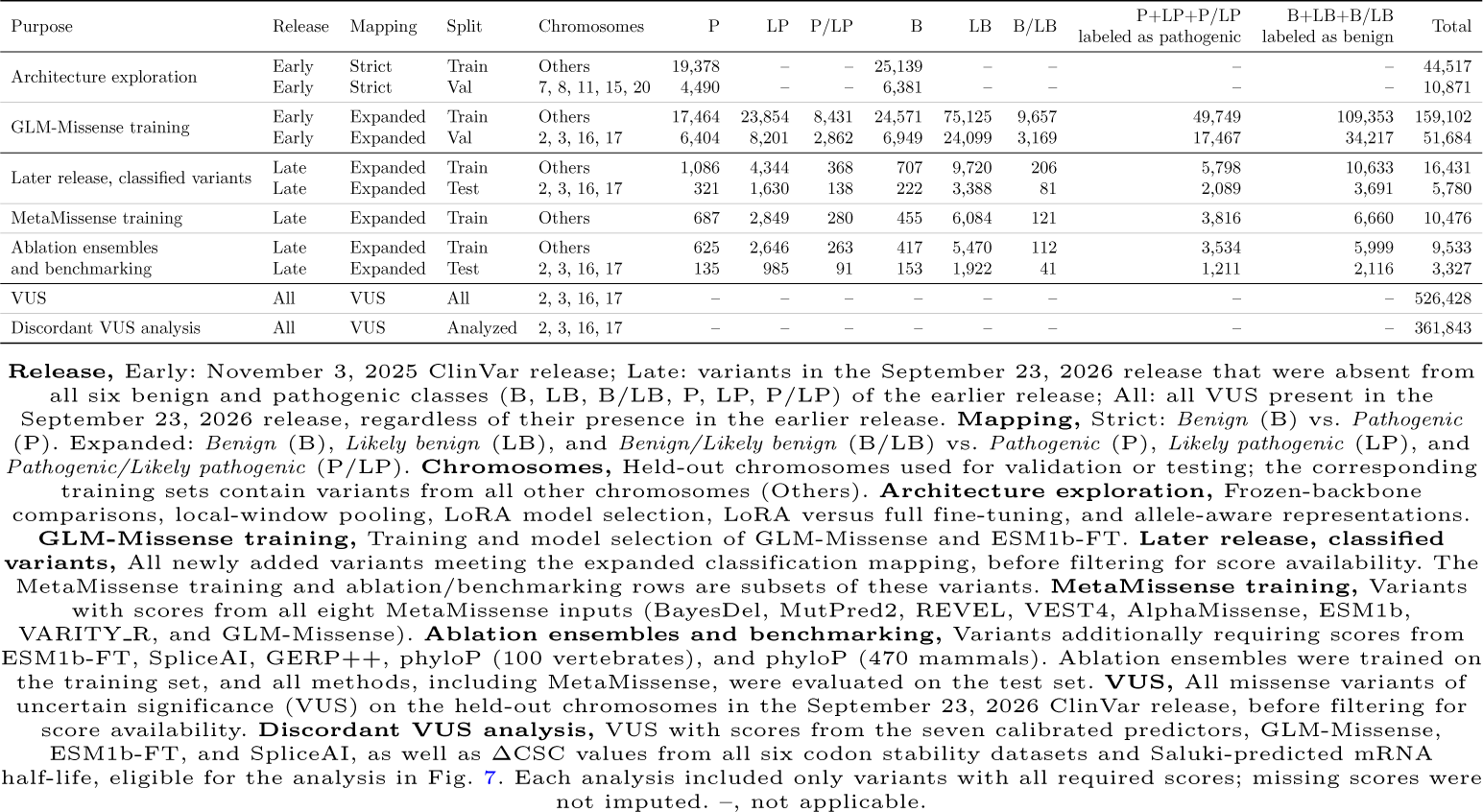
ClinVar sets used in this study. Variant counts by analysis, data release, classification mapping, and chromosome-based split.

#### Earlier-release sets

The earlier-release set was used in two settings, each with its own label mapping and held-out chromosomes.

##### Architecture exploration

To compare backbones, embedding strategies, classifier heads, adaptation regimes, and allele-aware inputs, we used earlier-release variants under the strict mapping. Chromosomes 7, 8, 11, 15, and 20 formed the validation set (*N* = 10,871), and all other chromosomes formed the training set (*N* = 44,517). For LoRA model selection, we used a 5,000-variant subset of the training set and a 1,250-variant subset of the validation set. The combined subset was repartitioned into five chromosome-exclusive folds for cross-validation. Only design choices and hyperpa-rameters were carried forward from these experiments; no trained model weights were reused.

##### GLM-Missense training

GLM-Missense and ESM1b-FT were trained on earlier-release variants under the expanded mapping. Chromosomes 2, 3, 16, and 17 formed the validation set (*N* = 51,684), and all other chromosomes formed the training set (*N* = 159,102). The validation set was used only for model selection.

#### Later-release sets

The later-release set was used to train the ensembles and to evaluate all methods. It was divided by the held-out chromosomes of GLM-Missense training into a training set (chromosomes other than 2, 3, 16, and 17; *N* = 16,431) and a test set (chromosomes 2, 3, 16, and 17; *N* = 5,780), both under the expanded mapping. No variant on the test chromosomes was used to train GLM-Missense, ESM1b-FT, or any ensemble.

Not every predictor scores every variant, and we did not impute missing scores, so that imputed values could not create artificial disagreement between methods. Each analysis therefore used only the variants with every score it required. MetaMissense was trained on the 10,476 training variants (3,816 labeled as pathogenic and 6,660 labeled as benign) with all eight of its inputs: the seven calibrated predictors and GLM-Missense. The ablation ensembles and all method comparisons used one shared subset: variants that also had ESM1b-FT, SpliceAI, GERP++, and phyloP scores. This subset comprised 9,533 training variants and 3,327 test variants (1,211 labeled as pathogenic and 2,116 labeled as benign). All ablation ensembles were trained on these 9,533 variants. All methods, MetaMissense included, were evaluated on the same 3,327 test variants.

#### VUS

For the discordant VUS analysis, we retrieved all missense variants classified as VUS in the September 23, 2026 ClinVar release on the held-out chromosomes (2, 3, 16, and 17), without excluding variants present in the earlier release. Of the 526,428, 365,724 had scores from the seven calibrated predictors, GLM-Missense, and ESM1b-FT. Of these, 361,843 also had SpliceAI, ΔCSC, and Saluki scores and were eligible for the discordant VUS analysis.

### 5.2 Backbone-specific input sequence construction

All sequence inputs were centered on the candidate missense variant so that the altered position occupied a fixed location after tokenization. Depending on the experiment, the model received either the alternate sequence alone or paired reference and alternate sequences processed separately and combined only at the classifier input. This design enabled consistent extraction of variant-position and local-context embeddings across backbones while allowing each model to operate at its native tokenization scheme and context length.

#### Nucleotide Transformer v2 (NT-2)

For each variant, we extracted an 11,999-bp sequence centered on the variant position and tokenized it using 6-mer encoding with stride 6. This yielded 2,000 tokens, with the variant located at token index 1,000. We used the pretrained Nucleotide Transformer v2 (NT-2, 500M parameters), a BERT-style model trained by masked language modeling on 850 genomes from diverse species Dalla-Torre et al. (2025). NT-2 produces a 1,024-dimensional embedding per token. For the matched NT-2 versus NT-1 comparison, we also evaluated NT-2 on a 5,999-bp input window.

### Nucleotide Transformer v1 (NT-1)

NT-1 is a 500M-parameter model with a similar architecture to NT-2 but pretrained on human genomes only. For NT-1, we extracted a 5,999-bp sequence centered on the variant and applied the same 6-mer tokenization strategy as NT-2, yielding 1,000 tokens with the variant at token index 500. NT-1 produces a 1,280-dimensional embedding per token.

#### Caduceus-PS

For Caduceus-PS, we extracted a 29,999-bp sequence centered on the variant. Caduceus-PS is a bi-directional Mamba-based DNA language model with reverse-complement equivariance achieved by parameter sharing Schiff et al. (2024). Unlike NT-1 and NT-2, it operates at single-nucleotide resolution. The resulting sequence therefore contains 29,999 tokens, with the variant at nucleotide index 15,000. Caduceus-PS produces a 512-dimensional embedding per token. Although Caduceus models were pretrained with substantially longer contexts (up to 131 kb) Schiff et al. (2024), we limited inputs to ∼30 kb to fit our available GPU memory while still capturing distal sequence context.

### 5.3 Embedding strategies, allele-aware inputs, classifier heads, and adaptation regimes

We systematically explored four model components: embedding aggregation strategy, allele-aware input representation, classifier head, and training adaptation regime.

#### Embedding aggregation strategies

For frozen-backbone and LoRA experiments, we compared several ways to derive the classifier input from backbone hidden states. *Variant-position embedding* extracts the embedding at the variant position (i.e., 1,024-d at token index 1,000 for NT-2, 1,280-d at token index 500 for NT-1, 512-d at token index 15,000 for Caduceus-PS; see Fig. S1). *Mean pooling* averages hidden states across all tokens in the input sequence, with attention masking for proper handling of padding tokens. For Caduceus-PS with CNN and Transformer heads, we applied a *downsampled sequence* representation in which the full 30-kb hidden-state sequence was reduced to 2,048 positions by adaptive average pooling, thereby reducing memory requirements.

#### Local-window pooling

To test whether pooling over a narrow neighborhood around the variant could recover local context without full-sequence dilution, we evaluated local-window mean pooling in the frozen NT-2 with CNN setting. Hidden states were mean-pooled over symmetric windows of ±8, ±16, or ±32 tokens around the variant position. With 6-mer tokenization, these correspond to ±48, ±96, and ±192 nucleotides, respectively.

#### Allele-aware input representations

Most frozen-backbone and LoRA experiments used only the variant-position embedding from the alternate sequence (*alt*). In the final input-representation comparison, we also evaluated *ref-alt*, which con-catenates the reference and alternate variant-position embeddings, and *ref-alt-diff*, which concatenates the reference embedding, alternate embedding, and their element-wise difference. For NT-2, these inputs have dimensions 1,024 (*alt*), 2,048 (*ref-alt*), and 3,072 (*ref-alt-diff*). To separate the effect of the allelic contrast from that of input dimension, we also evaluated an equal-dimension control, *alt-alt-alt*, which concatenates three copies of the alternate embeddings (3,072 dimensions).

#### Classifier heads

We evaluated three lightweight classifier heads: a multilayer perceptron (MLP), a convolutional neural network (CNN), and a shallow Transformer encoder. The MLP used either two hidden layers with dimensions 512 and 256, or three hidden layers with dimensions 512, 256, and 128, followed by a single-output classification layer. ReLU activations and 10% dropout were applied between hidden layers. The CNN head used two one-dimensional convolutional layers that projected the input hidden dimension to 256 and then to 128 channels, with kernel size 3, stride 1, padding 1, batch normalization, ReLU activation, and 10% dropout. The Transformer head used a two-layer Transformer encoder with hidden dimension 128, two attention heads, feed-forward dimension 512, and 10% dropout, followed by a linear classification layer. See Fig. S1 for details.

#### Training adaptation regimes

We evaluated three adaptation regimes. In the *frozen-backbone* setting, the pretrained backbone was fixed, and only the classifier head was trained. In the *full fine-tuning* setting, 100% of NT-2 parameters were updated. In the *LoRA* setting, the pretrained backbone remained fixed and only low-rank adaptation matrices (*W* = *W*_0_ + *BA* where *B* ∈ R*^d×r^*, *A* ∈ R*^r×d^*, *r* ≪ *d*) were trained Hu et al. (2022). We applied LoRA to the query and value projections in all attention layers and evaluated ranks *r* ∈ {8, 16, 32}, corresponding to approximately 0.19%, 0.38%, and 0.76% of total NT-2 parameters.

### 5.4 Optimization and training settings

Across all fine-tuning experiments, we used AdamW with *ɛ* = 10*^−^*^8^, *β*_1_ = 0.9, *β*_2_ = 0.999, and weight decay = 0.01, following the PyTorch defaults. We used a linear learning-rate schedule with warmup over the first 6% of training steps and linear decay to zero thereafter Liu et al. (2019). We optimized binary cross-entropy loss throughout. Unless otherwise noted, neural-network models were trained with early stopping using a patience of 2 epochs while monitoring validation AUROC. The comparison between LoRA and full fine-tuning used a stricter stopping rule. In that comparison, training stopped when either the train–validation gap exceeded 0.1 or the validation metric failed to improve for 2,000 consecutive steps.

Frozen-backbone experiments used batch size 32 and were run for up to 5,000 optimization steps. LoRA model selection experiments were performed on a reduced 5,000-variant subset obtained by stratified subsampling from each class. These experiments used batch size 4 and were run for up to 4,000 steps. Final GLM-Missense training and full fine-tuning used per-step batch sizes of 8 or 4, depending on GPU memory, together with gradient accumulation to maintain an effective batch size of 32. All experiments were run on 1–4 NVIDIA H100 GPUs (80 GB memory each).

Frozen-backbone experiments typically required ∼1–2 days per NT run and ∼4–6 hours per Caduceus run. LoRA training on the full earlier-release expanded training set required ∼3–5 days on one GPU. Full fine-tuning of NT-2 required ∼4 days on two GPUs.

### 5.5 Fine-tuning experimental procedures

#### Frozen-backbone comparisons

Frozen-backbone comparisons used the architecture-exploration training set (*N* = 44,517) and validation set (*N* = 10,871). We compared NT-2, NT-1, and Caduceus-PS across combinations of embedding strategies and classifier heads. The matched NT-2 versus NT-1 comparison used the same downstream configuration and a 5,999-bp NT-2 input window.

#### Local-window pooling

The local-window ablation was performed only in the frozen NT-2 setting with a CNN head. Using the architecture-exploration training and validation sets, we replaced full-sequence mean pooling with variant-centered local pooling over windows of ±8, ±16, and ±32 tokens.

#### LoRA model selection

LoRA model selection was performed on NT-2 using a 5,000-variant subset drawn by label-stratified random sampling from the architecture-exploration training set. Validation was performed on a 1,250-variant subset of the architecture-exploration validation set. We evaluated 54 configurations spanning two embedding strategies (variant-position and mean-pooling), three classifier heads (MLP, CNN, and Transformer), three LoRA ranks (*r* ∈ {8, 16, 32}), and three learning rates (1 × 10*^−^*^5^, 3 × 10*^−^*^5^, 5 × 10*^−^*^5^). The top ten configurations by validation performance were then re-evaluated by chromosome-exclusive 5-fold cross-validation on the combined 6,250-variant subset.

#### Full fine-tuning comparison

The LoRA versus full fine-tuning comparison used the architecture-exploration training and validation sets. We compared the top-ranked LoRA configuration against full fine-tuning of NT-2 at five learning rates from 1×10*^−^*^6^ to 1 × 10*^−^*^4^.

#### Allele-aware input comparison

For the allele-aware input comparison, we used the selected NT-2 LoRA configuration with an MLP classifier head. We com-pared four variant-position input representations: *alt*, *ref-alt*, *ref-alt-diff*, and the equal-dimensionality control *alt-alt-alt*.

### 5.6 GLM-Missense training and NT-2 zero-shot scoring

GLM-Missense used the NT-2 backbone, LoRA fine-tuning with rank *r* = 32 (scaling factor *α* = 64, LoRA dropout 0.1, applied to the query and value projections), learning rate 5 × 10*^−^*^5^, and the variant-position *ref-alt-diff* input representation. The reference and alternate sequences were encoded by the shared LoRA-adapted backbone; the variant-position embedding of each was passed through a shared MLP projector (1,024 to 512 to 256 dimensions), and the concatenation of the projected reference embedding, projected alternate embedding, and their difference (768 dimensions) was passed to a 2-layer MLP classifier head (768 to 512 to 256 to 1), with ReLU activations and 10% dropout. Training used the earlier-release expanded training set, and model selection used the earlier-release expanded validation set. The model was trained for 10,000 steps with an effective batch size of 32. Every 1,000 steps, performance was evaluated on 5,000 randomly drawn training variants and on a fixed subset of 5,000 validation variants drawn once across the validation chromosomes, and the checkpoint with the highest validation AUROC (step 8,000) was selected as GLM-Missense (Fig. S6).

For zero-shot scoring, we used the pretrained NT-2 backbone without task-specific fine-tuning. For each variant, we encoded matched reference and alternate sequences with the same 11,999-bp centered input window and 6-mer tokenization used for supervised NT-2 experiments. We then masked the k-mer token containing the variant position in the alternate sequence and computed a masked marginal log-likelihood ratio,

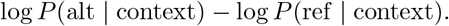

We transformed this quantity into a pathogenicity score by applying

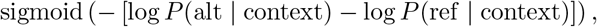

so that variants for which the alternate allele was less compatible with the pretrained model received higher pathogenicity scores. Zero-shot NT-2 scoring was evaluated on the later-release test set and compared directly with GLM-Missense.

### 5.7 ESM1b-FT, a protein-model control matched to GLM-Missense

ESM1b-FT was designed to differ from GLM-Missense mainly in its input. We replaced the NT-2 backbone of GLM-Missense with the ESM1b protein language model Rives et al. (2021) and kept all other components and settings. For each variant, we used the transcript that ClinVar used to name the variant. We built the reference and alternate protein sequences from the RefSeq protein sequence of this transcript. Vari-ants whose protein change could not be reconstructed or did not match this sequence were excluded. ESM1b accepts at most 1,022 residues. For longer proteins, we used a 1,022-residue window around the mutated residue, shifted inward at the protein ends. The reference and alternate sequences were encoded separately by ESM1b with LoRA (*r* = 32, *α* = 64, dropout 0.1, applied to the query and value projections). The embed-ding of the mutated residue was taken from each sequence, passed through a shared projector, and combined as *ref-alt-diff* before the MLP classifier head. Optimization followed GLM-Missense: AdamW with a learning rate of 5 × 10*^−^*^5^, 6% linear warmup followed by linear decay, and an effective batch size of 32. ESM1b-FT was trained on the same training set as GLM-Missense for 10,000 steps and evaluated every 1,000 steps on the same validation set. The checkpoint with the highest validation AUROC (step 3,000) was selected (Fig. S10).

### 5.8 Sources of external predictor scores and variant annotations

#### Comparator predictors

The comparator set comprised the seven missense variant impact predictors calibrated by ClinGen for the PP3 and BP4 criteria (Table 1 of Bergquist et al. (2025)): BayesDel without allele frequency, MutPred2, REVEL, VEST4, AlphaMissense, ESM1b, and VARITY R (Feng 2017; Pejaver et al. 2020; Ioan-nidis et al. 2016; Carter et al. 2013; Cheng et al. 2023; Brandes et al. 2023; Wu et al. 2021). Scores were obtained from the academic release of dbNSFP version 5.3.1a for GRCh38 (Liu et al. 2020) using the columns BayesDel noAF score, MutPred2 score, REVEL score, VEST4 score, AlphaMissense score, ESM1b score, and VARITY R LOO score. Variants were matched on chromosome, position, reference allele, and alternate allele. For transcript-specific scores, we used the score for the transcript that ClinVar used to name the variant.

#### Additional scores and variant annotations

Nucleotide conservation scores (GERP++ RS, phyloP100way vertebrate, and phyloP470way mammalian) were obtained from the same dbNSFP release (Davydov et al. 2010; Pollard et al. 2010; Christmas et al. 2023). gnomAD v4.1 joint allele frequencies (gnomAD4.1 joint AF) were collected from the same release. Splicing impact was taken from the raw SpliceAI SNV resource for GRCh38 (spliceai scores.raw.snv.hg38.vcf.gz) as the maximum of the four delta scores: acceptor gain, acceptor loss, donor gain, and donor loss (Jaganathan et al. 2019). We used raw rather than masked scores to retain predicted effects at both canonical and cryptic positions. As a null predictor, each variant received a random score drawn from a uniform distribution on [0, 1] (fixed random seed). All scores were oriented so that higher values indicate greater predicted pathogenicity; ESM1b scores were negated. Missing scores were not imputed: a variant without one of the scores used by an analysis was excluded from that analysis.

### 5.9 Pairwise concordance and partial correlation analyses

Each analysis included eight predictors: the seven established predictors and GLM-Missense for the ClinVar analysis (Fig. 4A, B); the same with ESM1b-FT in place of GLM-Missense for the protein-model control (Fig. S11A, B); and the seven established predictors with ESM1b-FT in place of ESM1b, plus GLM-Missense, for the *RAD51D* SGE analysis (Fig. 5A, B).

Pairwise concordance between predictor scores was measured by Kendall’s *τ* on the evaluation variants, and predictors were ordered by average-linkage hierarchical clustering on 1 − *τ* .

To measure the association between each predictor and the label that is not shared with the other predictors, we computed its partial Spearman correlation with the label. For each predictor (the focal predictor), the other seven predictors served as controls. The random score was included as a null reference: its partial correlation was computed with all eight predictors as controls. All scores and the binary label were rank-transformed. We then fit two ordinary least-squares regressions, each with an intercept, on the ranks of the controls: one with the ranks of the focal predictor as the outcome, and one with the ranks of the label as the outcome. The residuals of these regressions represent the parts of the focal predictor and of the label that the controls cannot explain, and the partial Spearman correlation is the correlation between these residuals. Larger positive values indicate that a predictor remains associated with the label after removing the variation it shares with the other predictors.

### 5.10 MetaMissense and ensemble ablation analyses

MetaMissense is an XGBoost ensemble (Chen and Guestrin 2016) of the seven calibrated predictors and GLM-Missense. It was trained on all later-release training variants with these eight scores (*n* = 10,476; 3,816 labeled as pathogenic; Table 2). Hyperparameters were selected by grid search, maximizing AUROC in chromosome-exclusive 5-fold cross-validation on these variants. The search space included n estimators ∈ {100, 300, 500}, max depth ∈ {3, 4, 5}, learning rate ∈ {0.01, 0.05, 0.1}, and subsample ∈ {0.8, 1.0}. The selected hyperparameters were n estimators = 500, max depth = 3, learning rate = 0.01, and subsample = 0.8 (cross-validated AUROC 0.984). The final MetaMissense model was trained with these hyperparameters on all 10,476 training variants and evaluated on the test variants with every score (*n* = 3,327).

All other ensembles were used to measure the information that a single predictor score adds to the others. Each such ablation analysis compared two ensembles that differed in only one input: one included the score of interest and the other did not. Everything else was the same. Both ensembles used the hyperparameters selected for MetaMissense, without further tuning, and both were trained once on the same later-release training variants (the 9,533 variants with every score). They were then evaluated on the same variants, either the ClinVar later-release test variants (*n* = 3,327) or the *RAD51D* variants (*n* = 1,819). Any difference in performance between the two ensembles therefore reflects the information carried by that one score of interest.

#### Ensemble ablation

To measure the contribution of each predictor, we compared a full ensemble with an ensemble trained identically without that predictor. For the ClinVar analyses (Fig. 4C), the full ensemble comprised the seven established predictors, GLM-Missense, and the random score; for the protein-model control (Fig. S11C), GLM-Missense was replaced by ESM1b-FT. For each predictor, we report the full ensemble minus the ensemble without that predictor, as ΔAUROC, ΔAUPRC, and log-loss reduction (the log-loss of the ensemble without the predictor minus that of the full ensemble), so that positive values indicate that including the predictor improves the ensemble. For each of GLM-Missense, GERP++, phyloP 100-way, and phyloP 470-way, we compared a base ensemble of the seven established predictors and the random score with ensembles in which each of these four scores was added separately (Fig. S9). Likewise, to measure the information that GLM-Missense adds beyond SpliceAI, we compared an ensemble with and without GLM-Missense, both containing the seven established predictors (with ESM1b-FT in place of ESM1b) and SpliceAI (Fig. 6B).

#### Standalone performance

We compared MetaMissense and each individual score on the ClinVar later-release test variants (*n* = 3,327) by AUROC, AUPRC, and log-loss reduction (Fig. S8). AUROC and AUPRC were computed directly from the scores. Log-loss requires predicted probabilities, which most individual scores do not provide. We therefore converted each score to a probability in two steps, both fitted on the later-release training variants (*n* = 9,533): a quantile-normal transformation based on the distribution of the score in these variants, followed by a logistic regression of the label on the transformed score. Both steps preserve the ranking of variants and therefore do not change AUROC or AUPRC. For MetaMissense, we used its own predicted probabilities. Log-loss reduction measured as standalone performance is the log-loss of a constant prediction equal to the proportion of pathogenic variants in the training variants, minus the log-loss of the score.

### 5.11 ClinVar label circularity analysis

ClinGen expert-panel classifications and the ACMG/AMP criteria applied to them were downloaded from the ClinGen Evidence Repository (https://erepo.clinicalgenome.org/evrepo/; accessed October 29, 2026). Classifications were matched to later-release variants by ClinVar Variation ID, and retracted classifications were excluded. We retained the variants with scores from all seven predictors: 176 pathogenic or likely pathogenic and 47 benign or likely benign, of 276 pathogenic or likely pathogenic and 111 benign or likely benign with expert-panel classifications. For each variant, we scored the applied criteria with the Bayesian point system: support-ing, 1 point; moderate, 2; strong, 4; very strong, 8; pathogenic, ≥ 10; likely pathogenic, 6–9; VUS, 0–5; likely benign, −1 to −6; benign, ≤ −7 (Tavtigian et al. 2020). We scored each classification with all criteria but without PP3 and BP4. A variant was considered dependent on PP3 or BP4 if, without these criteria, it no longer met the threshold of its original classification. We computed each predictor’s AUROC before and after removing the dependent variants, and the change in AUROC between the two.

### 5.12 *RAD51D* saturation genome editing data

*RAD51D* saturation genome editing data were obtained from Casadei et al. (2026).In this study, nearly all possible single-nucleotide variants in the targeted regions of *RAD51D* were introduced into the endogenous locus of HAP1 cells by CRISPR–Cas9-mediated homology-directed repair. The effect of each variant on cell fitness was measured by deep sequencing over time. Casadei et al. assigned each variant a functional classification and, for a subset of variants, an RNA score. The RNA score is the log_2_ ratio of the variant’s frequency in RNA to its frequency in genomic DNA from the same cells. We retained missense single-nucleotide variants classified as functionally abnormal (label 1; *n* = 227) or functionally normal (label 0; *n* = 1,592) with scores from every method, and excluded variants with other classifications. Variants were annotated relative to transcript ENST00000345365.11. *RAD51D* lies on chromosome 17, a held-out chromosome, so no *RAD51D* variant was used to train GLM-Missense, ESM1b-FT, or any ensemble. The ensembles used in the SGE analyses were trained on the ClinVar later-release training variants (*n* = 9,533), with ESM1b-FT in place of ESM1b, and then directly applied to the *RAD51D* variants.

### 5.13 Characterization of resolved variants

For each predictor, we classified each variant as pathogenic, benign, or neither, based on its score. For the calibrated predictors, we used the supporting-strength intervals of Bergquist et al. (2025) (Table S3). A score at or above the lower bound of PP3 supporting was called pathogenic, and a score at or below the upper bound of BP4 supporting was called benign. Scores between these bounds received no call. GLM-Missense and ESM1b-FT were called pathogenic at a score of at least 0.5 and benign otherwise.

Within each label class, the variants *resolved* by a focal model (GLM-Missense or ESM1b-FT) were those that it classified correctly while at least five of the seven reference predictors did not. The background set consisted of all other variants of the same class. In the ClinVar later-release test set, we compared the SpliceAI scores of resolved and background variants; in the RAD51D SGE data, we compared their RNA scores. Each comparison used a two-sided Mann–Whitney U test within each label class.

### 5.14 Discordant VUS analysis

Of the 526,428 ClinVar missense VUS on chromosomes 2, 3, 16, and 17, we retained the 361,843 with scores from GLM-Missense, ESM1b-FT, and the seven calibrated predictors, and with all three features described below (Table 2). Cases and controls were selected from these VUS.

As in the resolved-variant analysis, each calibrated predictor classified each VUS as pathogenic, benign, or neither, using its ClinGen supporting-strength thresholds (Table S3). When we counted how many predictors made a given call, a predictor that classified a VUS as neither was counted toward neither call. For each focal model (GLM-Missense or ESM1b-FT) and direction, a *case* was a VUS that the model called confidently while at least five of the seven calibrated predictors made the opposite call. Confident calls were a score of at least 0.8 (pathogenic; *to pathogenic* direction) or at most 0.2 (benign; *to benign* direction). Each case was matched to one *control* VUS, without replacement. A VUS was eligible as a control for a case if it was in the same gene, if the same number of calibrated predictors made the same call on it as on the case, and if the focal model called it confidently in agreement with those predictors. Among the eligible VUS not yet used as controls, we chose the one nearest to the case in genomic position, breaking ties at random. Cases were matched with controls in order of decreasing confidence of the focal model, and cases without an eligible control were dropped.

We compared cases with their matched controls for three nucleotide-level features. Predicted splicing disruption was the maximum SpliceAI delta score. Codon stability was measured as ΔCSC, the codon stability coefficient (CSC) of the alternate codon minus that of the reference codon (Wu et al. 2019). CSC values differ between cell types and assays. We therefore averaged ΔCSC over the six human CSC sets of Wu et al. (2019): endogenous transcripts in HEK293T, HeLa, and RPE cells; ORFeome libraries in HEK293T and K562 cells; and SLAM-seq in K562 cells. Codons were determined on the transcript on which ClinVar described the variant, using Ensembl release 115 annotations. Predicted mRNA stability was the change in Saluki-predicted mRNA half-life between the alternate and reference mature transcripts (Agarwal and Kelley 2022). Mature transcripts were constructed from the same transcript and Ensembl release. Predictions were taken from the human output of each of the 50 released Saluki models (10 cross-validation folds, 5 replicate training runs each) and averaged. Transcripts longer than 12,288 nt kept their last 12,288 nt, as in the original model; variants in the removed 5*^′^* region were not scored. Cases were compared with their own controls by two-sided Wilcoxon signed-rank tests on the paired differences. *P* values were adjusted by the Benjamini–Hochberg procedure over the six tests shown for each model (three features, two directions).

### 5.15 Evaluation metrics and statistical tests

The primary performance metrics were AUROC and AUPRC. For ensemble com-parisons, we also report the reduction in log-loss, which measures improvement in predicted probabilities rather than in ranking alone. For single train/validation runs, model selection used validation AUROC unless otherwise noted. For 5-fold cross-validation of LoRA configurations, we report the mean and standard error across folds.

Confidence intervals for the standalone performance of each score (Fig. S8), partial correlations (Figs. 4B, 5B, and S11B), ensemble comparisons (Figs. 4C, 5C, 6B, S9, and S11C), and the change in AUROC in the circularity analysis (Fig. S12C) were obtained from 1,000 class-stratified bootstrap resamples of the evaluation variants. For each comparison of two models, both were evaluated on the same resampled variants and their difference was computed within each resample, so that comparisons were paired. In the circularity analysis, each resample was drawn from all variants with expert-panel classifications. On each resample, we computed each predictor’s AUROC twice: on all resampled variants, and on only those whose labels did not depend on PP3 or BP4. The change in AUROC was the difference between the two. Resamples in which this subset contained variants of only one class, so that its AUROC could not be computed, were discarded. We report 95% percentile confidence intervals and two-sided bootstrap *p* values, computed as twice the smaller fraction of resampled statistics on either side of zero (minimum 1*/*1,000). Within each panel, *p* values were adjusted for multiple testing by the Benjamini–Hochberg procedure Benjamini and Hochberg (1995) (Figs. 4B–C, 5B–C, S9, S11B–C, and S12C). In Fig. S8, a score was considered not different from the top-ranked score if the unadjusted two-sided bootstrap *p* of their difference was ≥ 0.05. The comparison of GLM-Missense conditional on SpliceAI (Fig. 6B) was a single pre-specified test and was not adjusted. Resolved and background variants were compared with two-sided Mann–Whitney *U* tests, one pre-specified comparison per label class, without adjustment (Figs. 6A and S13). Discordant VUS were compared with their matched controls by two-sided Wilcoxon signed-rank tests, with Benjamini–Hochberg adjustment over the six tests shown for each model (three features, two directions; Figs. 7 and S14).

## Declarations

### Ethics approval and consent to participate

Not applicable.

### Consent for publication

Not applicable.

## Availability of data and materials

All code used to develop GLM-Missense and MetaMissense, together with scripts for the analyses reported in this study, is available at https://github.com/yaqisu/ GLM-Missense MetaMissense. ClinVar data are available from the NCBI ClinVar FTP site at https://ftp.ncbi.nlm.nih.gov/pub/clinvar/. dbNSFP data are available at https://www.dbnsfp.org/. Precomputed SpliceAI scores are available through Illu-mina BaseSpace at https://basespace.illumina.com/analyses/194103939/files; access requires registration for an Illumina account. The *RAD51D* saturation genome editing data are available from Casadei et al. Casadei et al. (2026). ClinGen expert-panel classifications are available from the ClinGen Evidence Repository at https://erepo.clinicalgenome.org/evrepo/. Human codon stability coefficients are available from Wu et al. Wu et al. (2019). The Saluki models are available from Zenodo (https://doi.org/10.5281/zenodo.6326409).

## Authors’ contributions

Y.S. and Y.L. designed the project, conducted the experiments, and analyzed the results. Y.L. and Y.S. wrote and reviewed the manuscript.

## Acknowledgements

The authors thank Dr. Steven E. Brenner for supporting the authors through the Brenner Lab, and for advice on related projects that inspired parts of the analysis. They thank Garv Goswami for his initial input on the Caduceus fine-tuning analysis, and the instructors and classmates of the UC Berkeley CS182/282A class (2025 Fall) for their early feedback and suggestions on the fine-tuning analysis.

## 6 Supplementary Figures and Tables

**Fig. S1.**
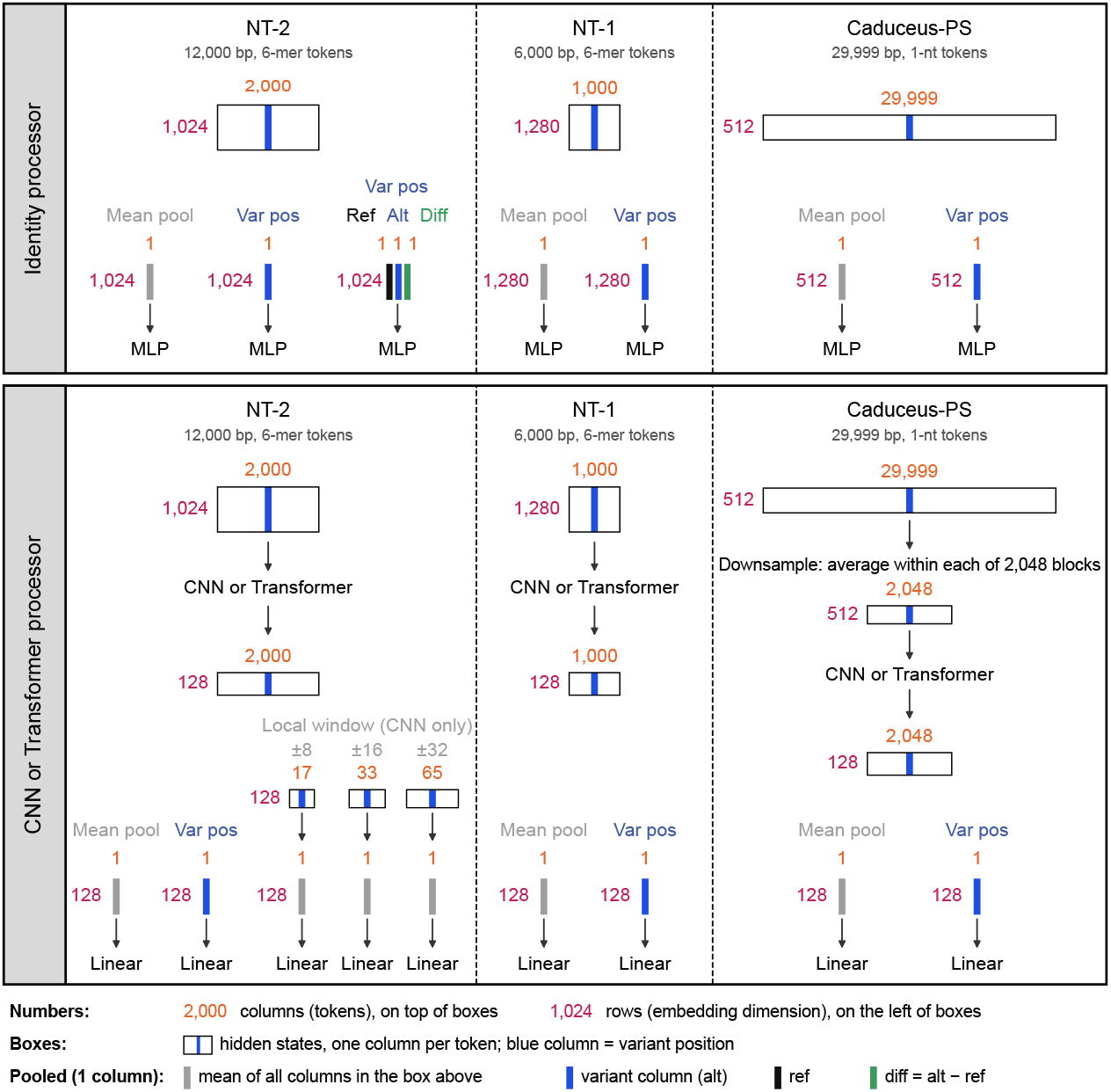
Overview of the backbone, aggregation, and classifier configurations explored for adapting genomic language models to clinical missense variant impact prediction. Three backbones were compared: NT-2 (12,000-bp input, 6-mer tokens; 2,000 tokens *×* 1,024 dimensions), NT-1 (6,000-bp input, 6-mer tokens; 1,000 *×* 1,280), and Caduceus-PS (29,999-bp input, single-nucleotide tokens; 29,999 *×* 512). (Top) Identity processor: the backbone hidden states are aggregated directly, either by mean pooling over all tokens or by taking the embedding at the variant position, and passed to an MLP. For NT-2, the *ref-alt-diff* representation concatenates the variant-position embeddings of the reference and alternate sequences and their difference (alt *−* ref); the *ref-alt* representation (not shown) concatenates the reference and alternate embeddings without the difference. (Bottom) CNN or Transformer processor: the hidden-state sequence is first processed by a CNN or a shallow Transformer, which reduces the embedding dimension to 128. It is then aggregated by mean pooling, by taking the variant position, or, for NT-2 with the CNN processor, by mean pool-ing over a local window of *±*8, *±*16, or *±*32 tokens around the variant (17, 33, or 65 tokens), before a linear output layer. For Caduceus-PS, the hidden-state sequence was first downsampled to 2,048 positions by averaging within blocks. Rectangles represent hidden-state matrices, with one column per token and the variant position marked in blue; orange numbers above each rectangle give the number of columns (tokens), and magenta numbers to its left give the number of rows (embedding dimension). Bars represent single-column embeddings passed to the classifier: gray, the mean of all columns in the rectangle above; blue, the variant-position column of the alternate sequence; black, the variant-position column of the reference sequence; green, the difference between alternate and reference (alt *−* ref).

**Fig. S2.**
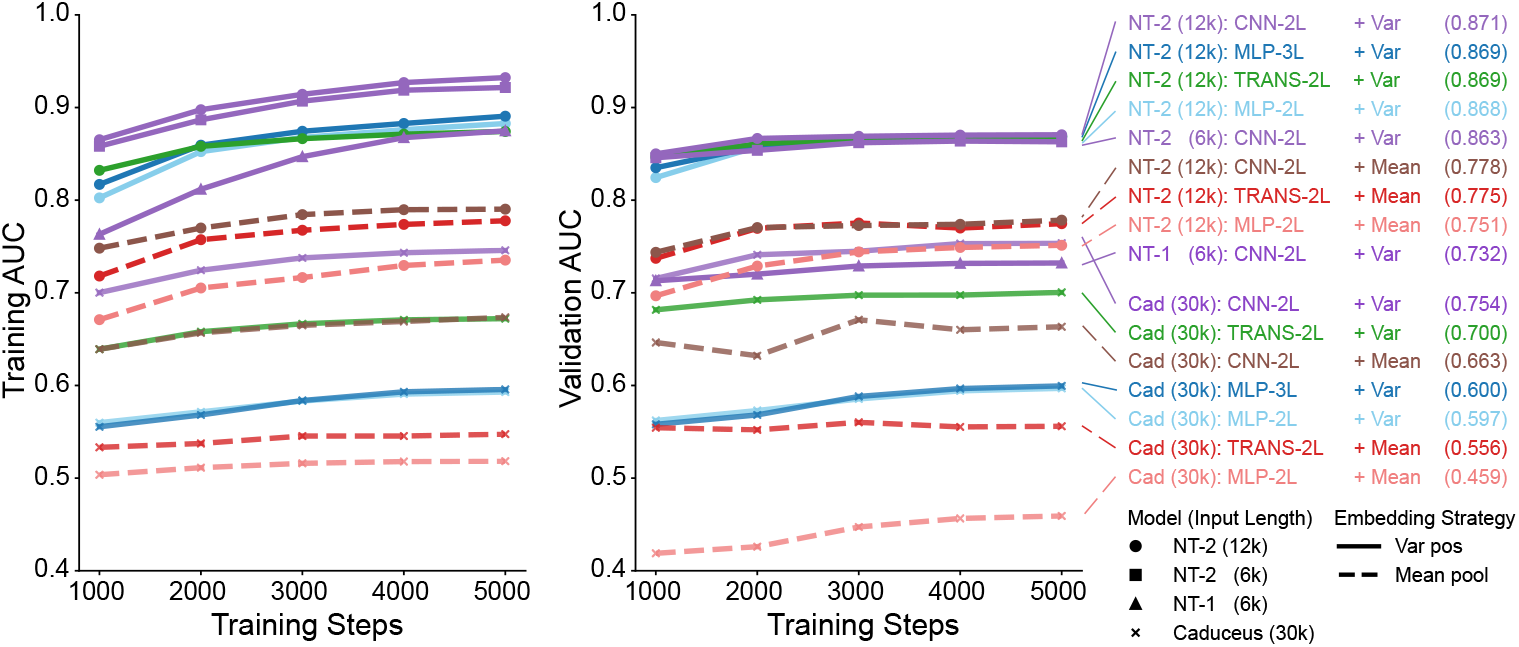
Training and validation curves for frozen-backbone comparison across genomic language model backbones, embedding strategies, and classifier heads. Training (left) and validation (right) AUC are shown over 5,000 training steps for combinations of three back-bone architectures: Nucleotide Transformer v2 (NT-2, 500M parameters), Nucleotide Transformer v1 (NT-1, 500M parameters), and Caduceus-PS (7.7M parameters), with four classifier heads: two-layer CNN (CNN-2L), three-layer MLP (MLP-3L), two-layer MLP (MLP-2L), and two-layer Transformer (TRANS-2L). Solid lines denote variant-position embeddings, while dashed lines denote mean-pooling embeddings. Marker shapes distinguish backbone models; line colors group experiments by backbone and embedding strategy. Numbers in parentheses in the legend indicate input sequence length for each experiment, not model size. Values in parentheses at the far right report the best validation AUC for each configuration.

**Fig. S3.**
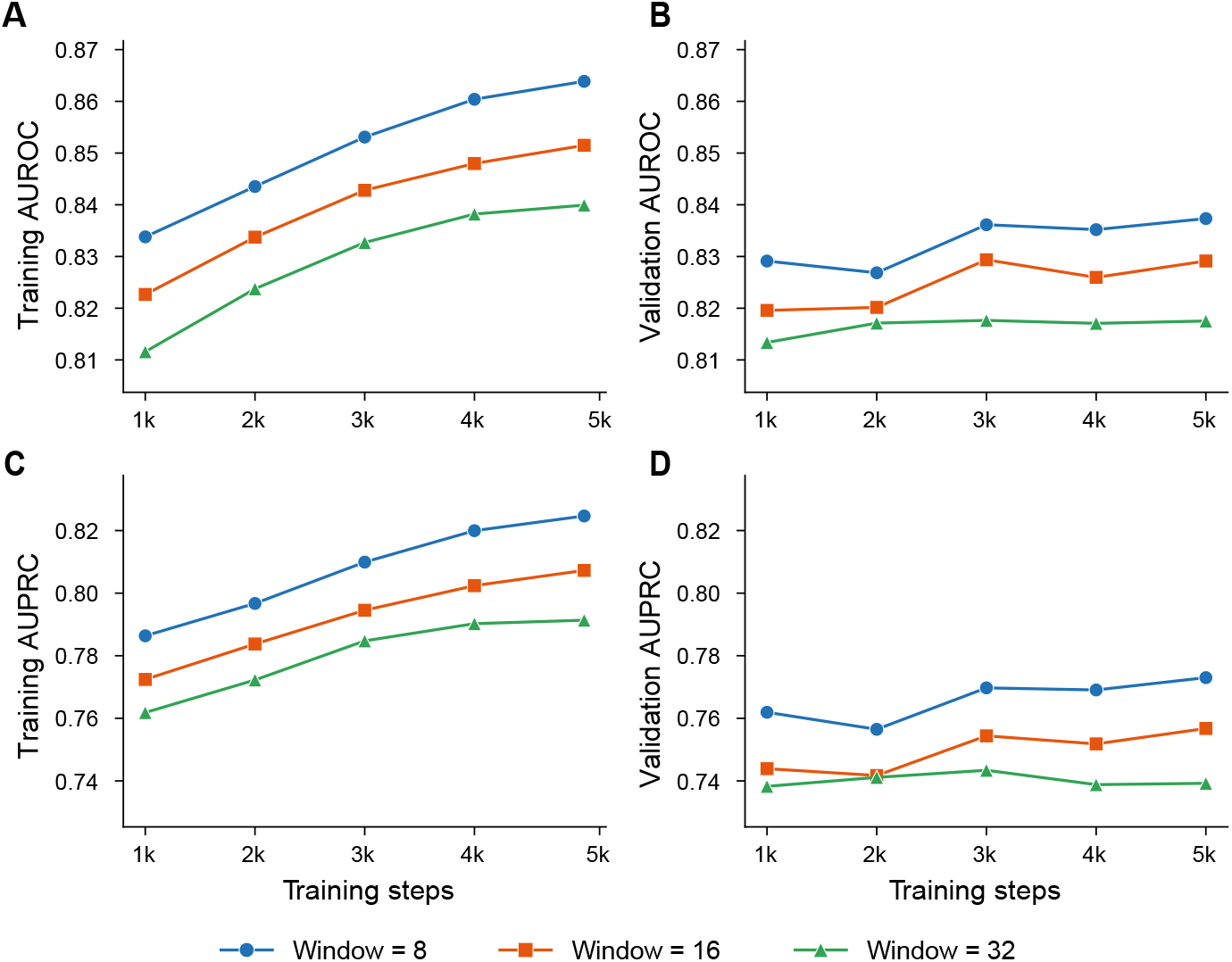
Local-window mean pooling performance across window sizes. Performance of frozen NT-2 with a CNN classifier head using mean pooling over *±*8, *±*16, or *±*32-token windows cen-tered on the variant. All models were trained on the earlier-release strict training set. (**A**) Training AUROC. (**B**) Validation AUROC. (**C**) Training AUPRC. (**D**) Validation AUPRC. Smaller windows performed better than larger windows, with the 8-token window giving the best validation perfor-mance among the local-window settings.

**Fig. S4.**
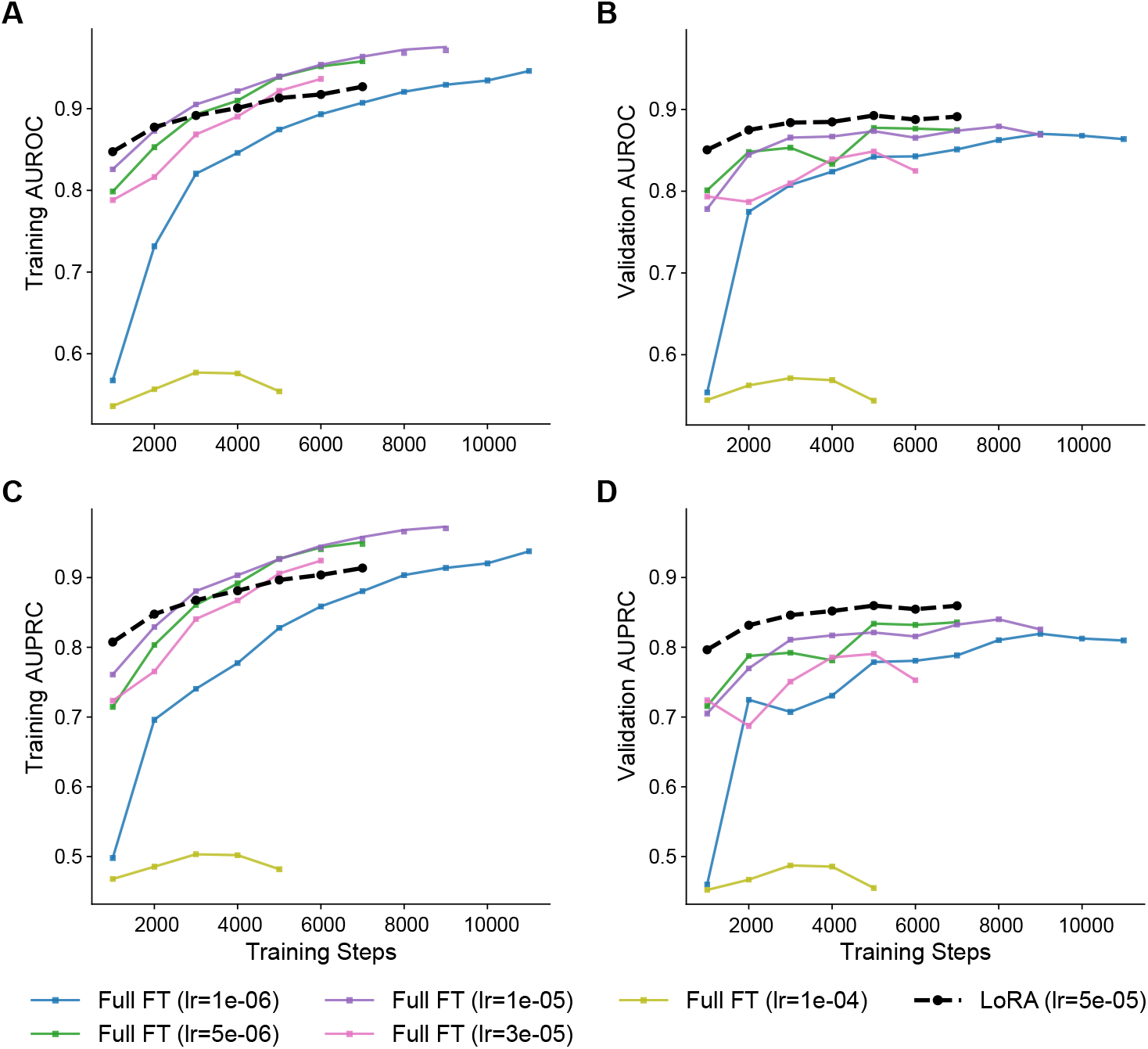
LoRA improves validation performance and generalization relative to full fine-tuning. Each panel shows one LoRA run (dashed black line) and five full fine-tuning runs of NT-2 at distinct learning rates. (**A**) Training AUROC. (**B**) Validation AUROC. (**C**) Training AUPRC. (**D**) Validation AUPRC. LoRA reaches the highest validation performance while maintaining a smaller train–validation gap.

**Fig. S5.**
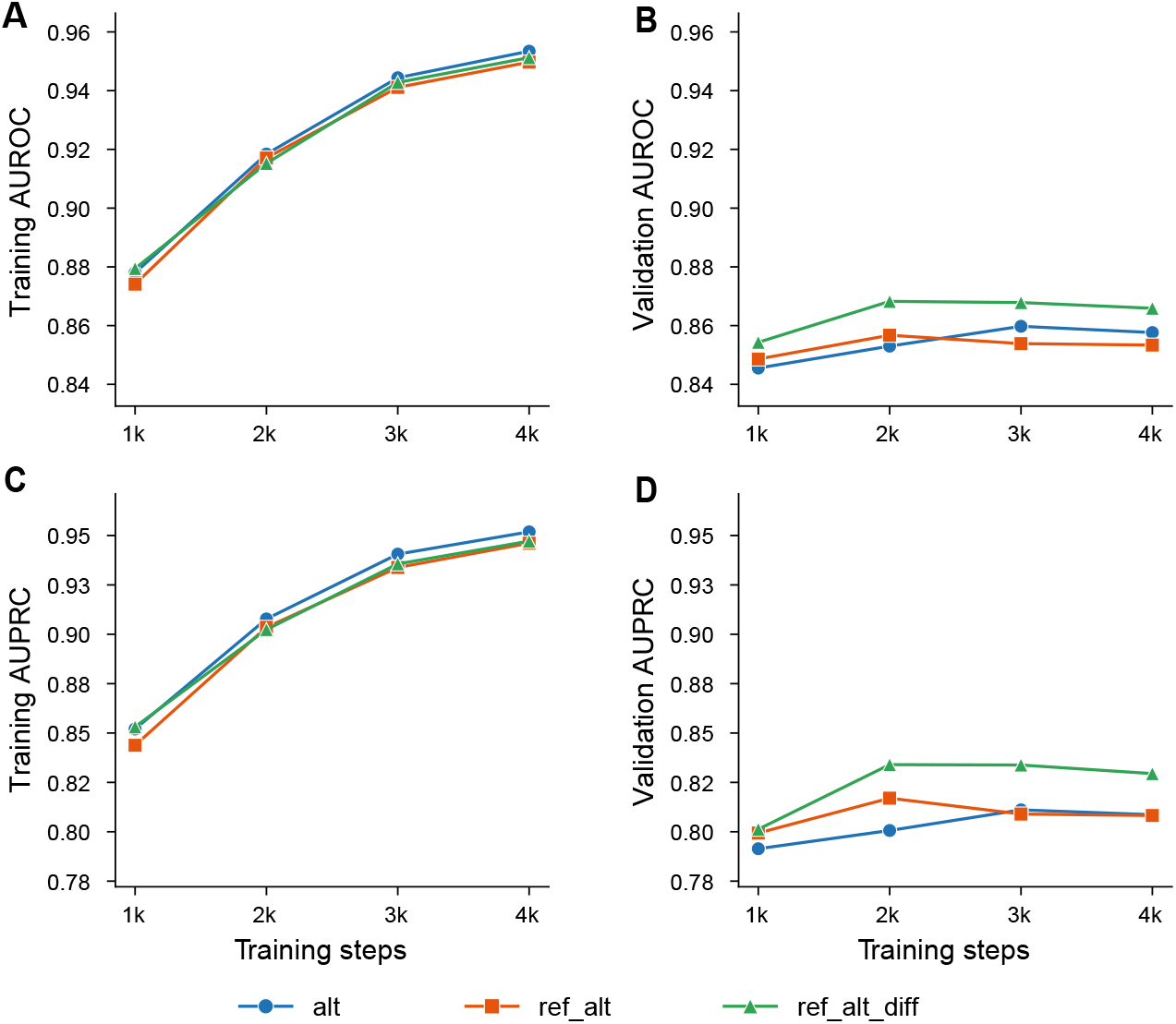
Allele-aware input representation performance across *alt*, *ref-alt*, and *ref-alt-diff*. Performance of NT-2 with LoRA rank 32 and an MLP classifier head trained on the 5k subset of the earlier-release strict training set. (**A**) Training AUROC. (**B**) Validation AUROC. (**C**) Training AUPRC. (**D**) Validation AUPRC. The *ref-alt-diff* representation gave the strongest validation per-formance, whereas *alt* and *ref-alt* performed similarly.

**Fig. S6.**
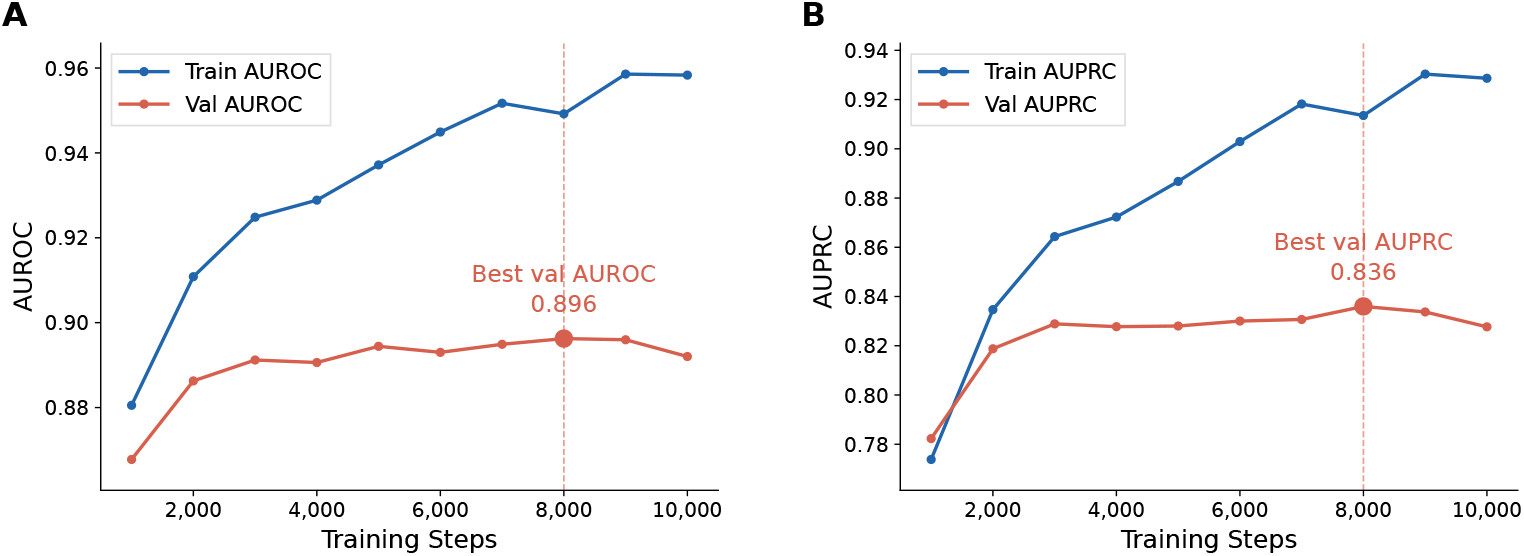
Training curves of the GLM-Missense model. Training and validation performance of the GLM-Missense configuration across optimization steps on the earlier-release expanded set. (**A**) AUROC. (**B**) AUPRC. At each checkpoint (every 1,000 steps), training performance was computed on 5,000 training variants drawn at random at that checkpoint, and validation performance on a fixed subset of 5,000 validation variants drawn once across the validation chromosomes (2, 3, 16, and 17). The vertical dashed line marks the step with the best validation metric (step 8,000), whose checkpoint was selected as GLM-Missense.

**Fig. S7.**
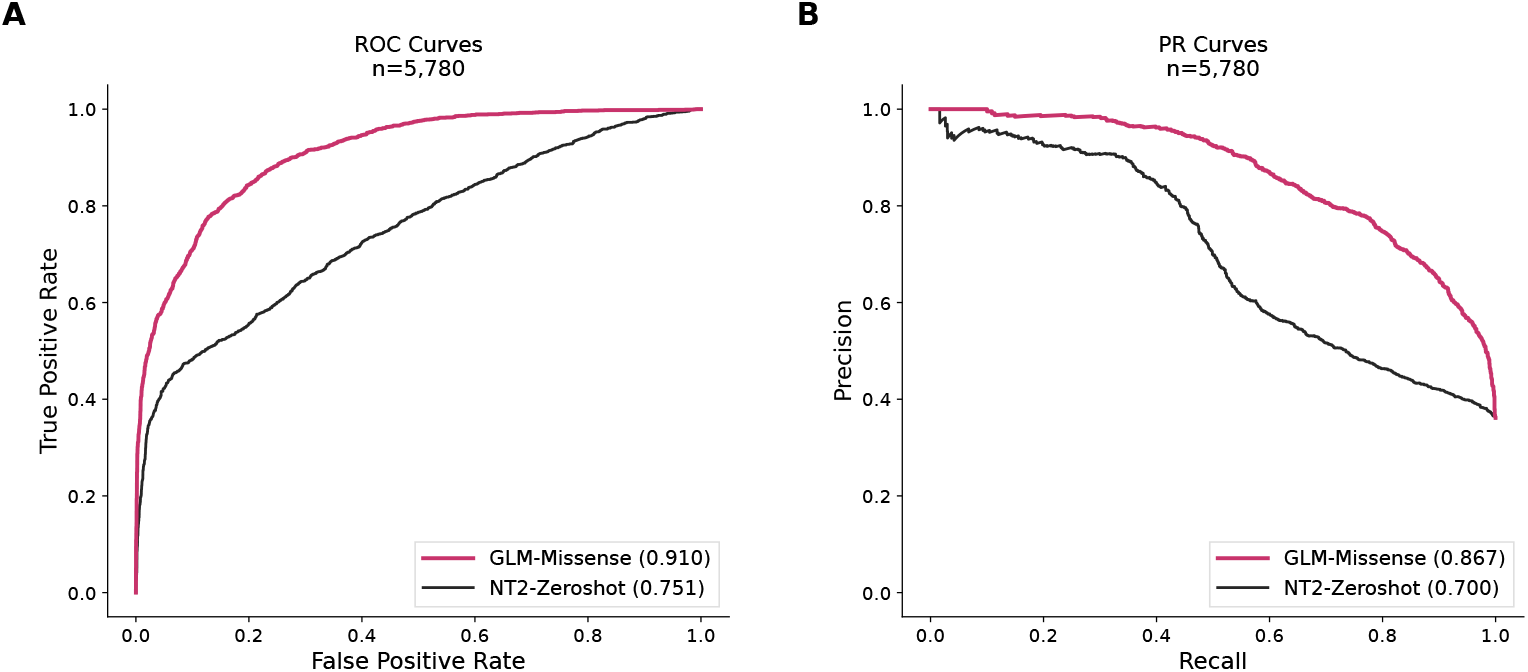
Fine-tuned GLM-Missense outperforms zero-shot NT-2 scoring. Comparison of GLM-Missense and zero-shot NT-2 on the later-release test set (*n* = 5,780). (**A**) ROC curves. (**B**) PR curves. GLM-Missense outperformed zero-shot NT-2 on both AUROC (0.910 versus 0.751) and AUPRC (0.867 versus 0.700).

**Fig. S8.**
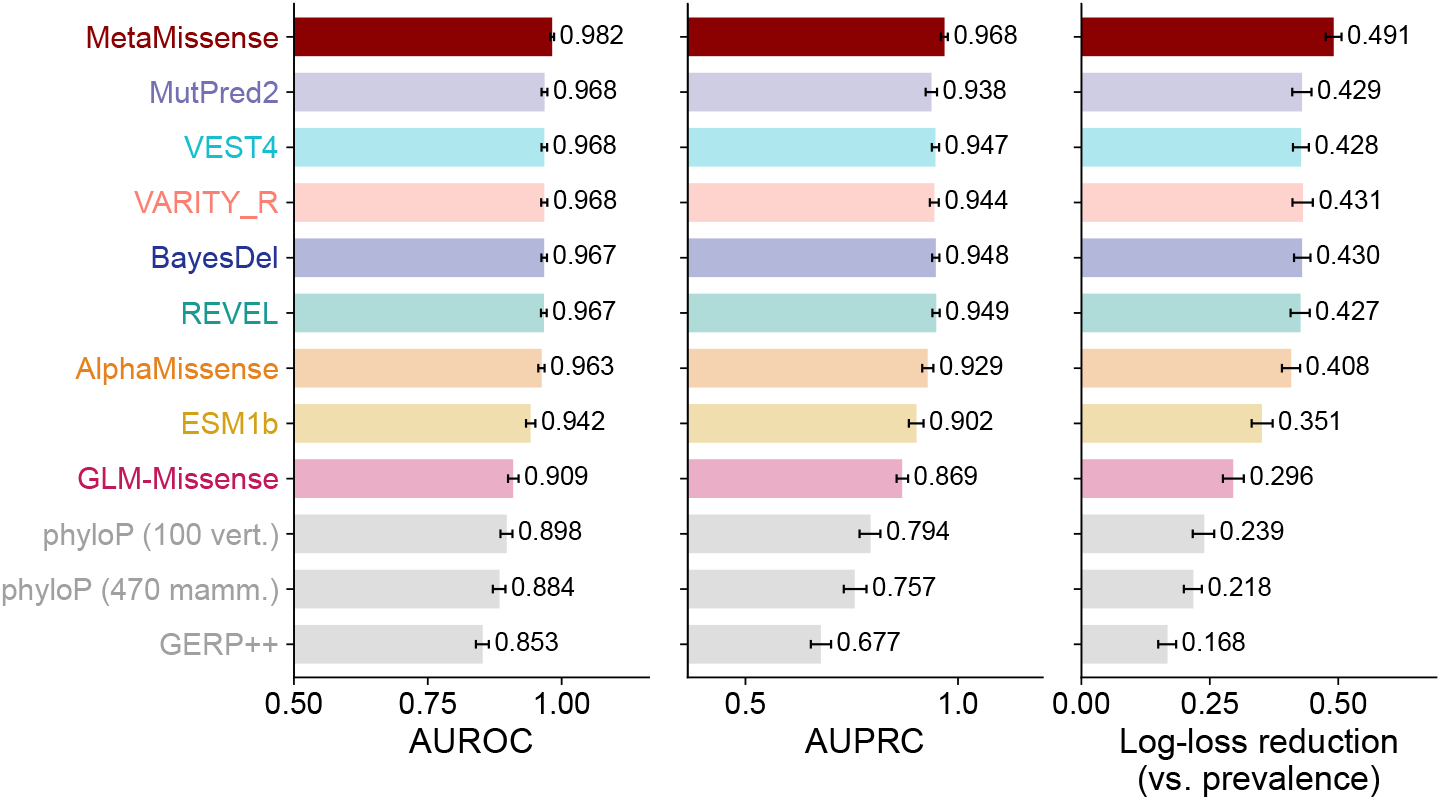
Standalone performance of MetaMissense and individual scores on held-out ClinVar variants. AUROC, AUPRC, and log-loss reduction relative to a constant prediction of the training-set prevalence on the ClinVar later-release test variants (*n* = 3,327; 1,211 labeled as pathogenic and 2,116 labeled as benign) for MetaMissense (the seven established predictors plus GLM-Missense), each of the seven comparator predictors, GLM-Missense, and the conservation scores phyloP 100-way vertebrate, phyloP 470-way mammalian, and GERP++. The log-loss of individual scores was computed after a quantile-normal transformation and logistic regression fitted on the later-release training set. Error bars denote 95% CIs from 1,000 class-stratified bootstrap resamples. Solid bars indicate scores not significantly different from the best-performing score (paired two-sided bootstrap *p ≥* 0.05); light bars indicate scores significantly worse than the best-performing score.

**Fig. S9.**
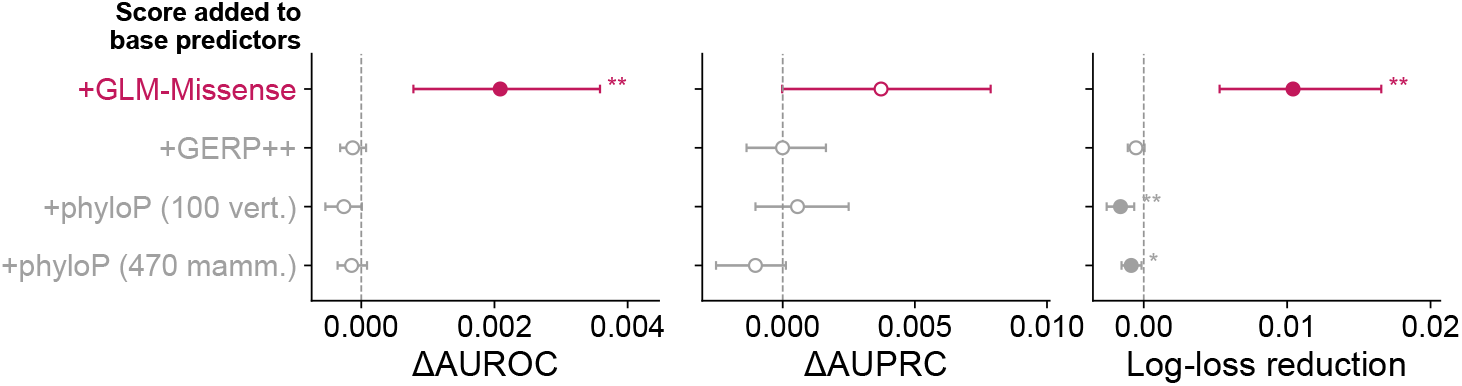
GLM-Missense, but not nucleotide-level conservation scores, improves an ensemble of the established predictors. Gain from adding each score (GLM-Missense, GERP++, phyloP 100-way vertebrate, or phyloP 470-way mammalian) to an ensemble of the seven established predictors and the random score, trained identically on the later-release training set (*n* = 9,533) and evaluated on the later-release test set (*n* = 3,327). Positive ΔAUROC, ΔAUPRC, and log-loss reduc-tion indicate that adding the score improves the ensemble. Error bars denote 95% CIs from 1,000 class-stratified paired bootstrap resamples. Filled symbols and stars denote Benjamini–Hochberg *q <* 0.05 within each panel: ^∗^ *q <* 0.05, ^∗∗^ *q <* 0.01; open symbols are not significant.

**Fig. S10.**
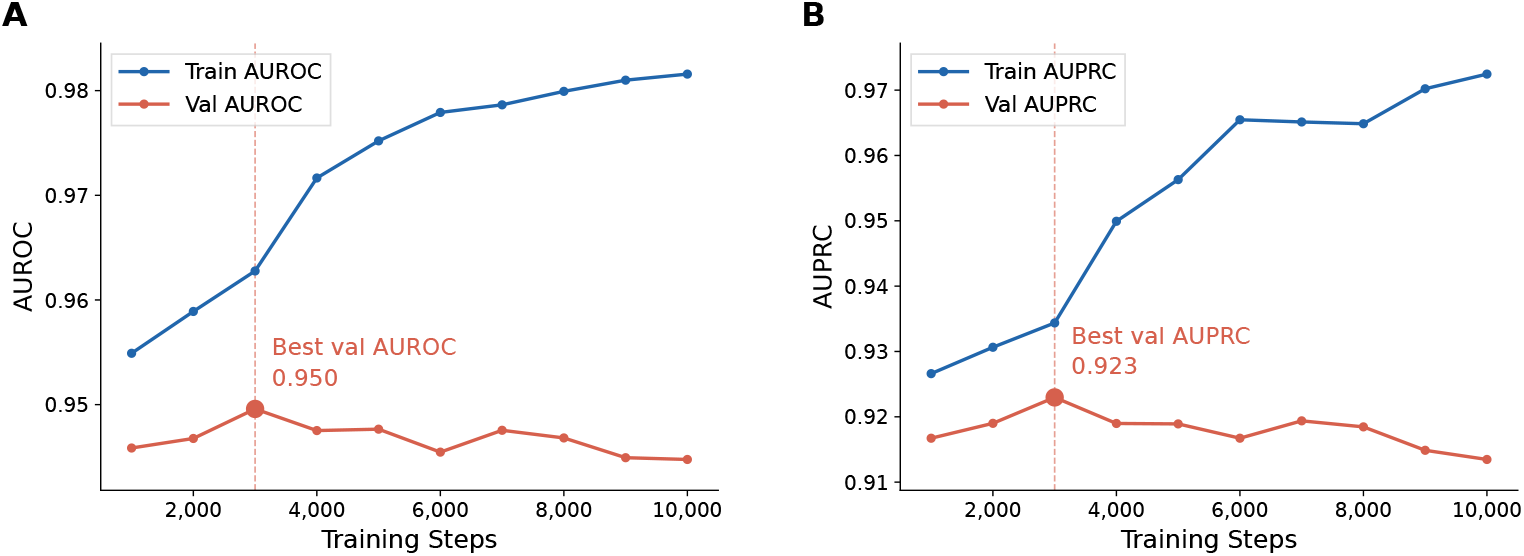
Training curves of the matched protein-model control ESM1b-FT. Training and validation performance of ESM1b fine-tuned with LoRA on the earlier-release expanded training set, using the same architecture (apart from the backbone), hyperparameters, and training procedure as GLM-Missense. (**A**) AUROC. (**B**) AUPRC. Training performance was computed on randomly drawn training variants and validation performance on the same validation subset as for GLM-Missense (Fig. S6). The vertical dashed line marks the step with the best validation metric (step 3,000), whose checkpoint was selected as ESM1b-FT.

**Fig. S11.**
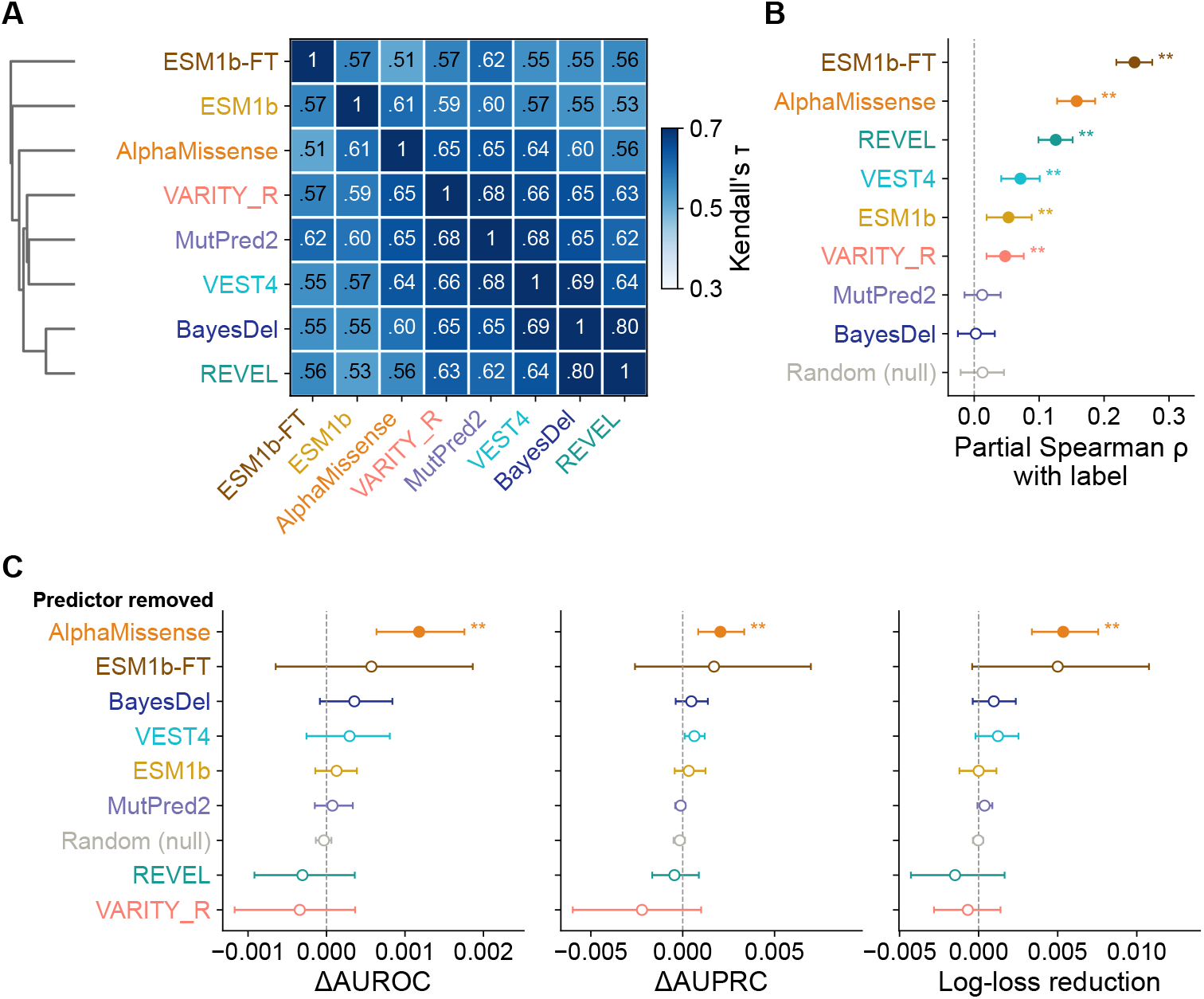
A protein language model fine-tuned on the same training variants as GLM-Missense does not contribute complementary signal to the ensemble. The analyses of Fig. 4, with ESM1b-FT in place of GLM-Missense, on the ClinVar later-release test variants (*n* = 3,327). (A) Pairwise Kendall’s *τ* between ESM1b-FT and the seven established predictors, with average-linkage hierarchical clustering based on 1 *− τ* . (B) Partial Spearman correlation between each predictor and the ClinVar label after controlling for the other seven predictor scores; Random (null) is controlled for all eight predictors. (C) Ensemble ablation analysis, as in Fig. 4C, for an ensemble of the seven established predictors, ESM1b-FT, and the random score, trained on the later-release training set (*n* = 9,533). Positive values indicate that including the predictor improves the ensemble. Error bars denote 95% CIs from 1,000 class-stratified bootstrap resamples, paired across rows. Filled symbols and stars denote Benjamini–Hochberg *q <* 0.05 within each panel: ^∗∗^ *q <* 0.01; open symbols are not significant.

**Fig. S12.**
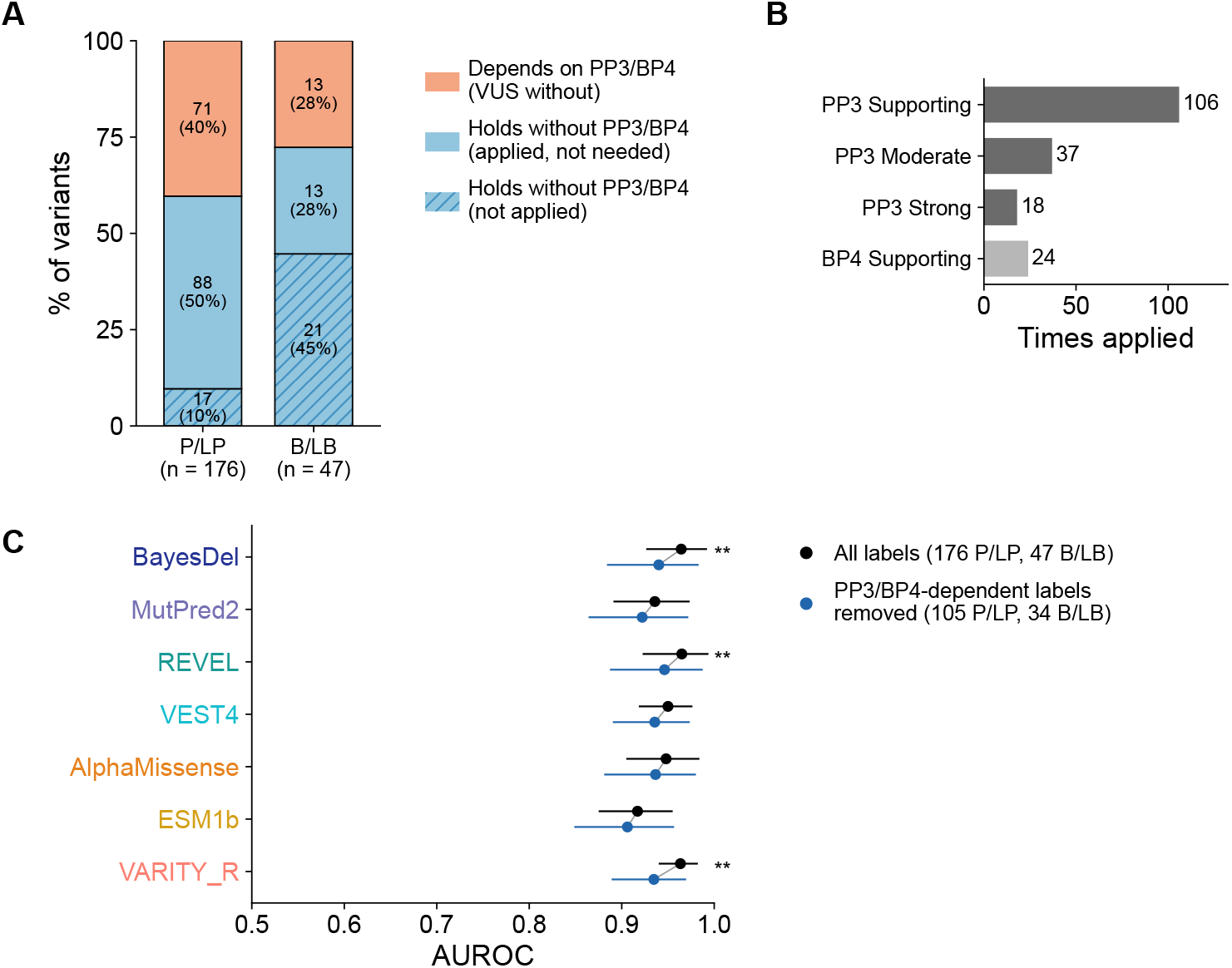
ClinVar labels partly depend on computational evidence, which inflates the performance of the established predictors. Later-release variants (all chromosomes) with Clin-Gen expert-panel classifications and scores from all seven established predictors (176 pathogenic or likely pathogenic, 47 benign or likely benign) were collected. Each classification was re-scored with the Bayesian point system without the computational criteria PP3 and BP4 Tavtigian et al. (2020). **(A)** Share of variants whose label holds without PP3 or BP4 (hatched, criteria not applied; solid blue, applied but not needed) or depends on them (orange; the variant would be a VUS without them). Numbers indicate variants and percentages. **(B)** Number of times PP3 and BP4 were applied, by strength. **(C)** AUROC of each predictor with all labels (black) and after removing labels that depend on PP3 or BP4 (blue; 105 pathogenic or likely pathogenic, 34 benign or likely benign). Error bars, 95% confidence intervals from 1,000 class-stratified bootstrap resamples; the reduced set was taken within each resample. Stars, Benjamini–Hochberg-adjusted *q* of the change in AUROC across the seven predictors (\*\**q <* 0.01).

**Fig. S13.**
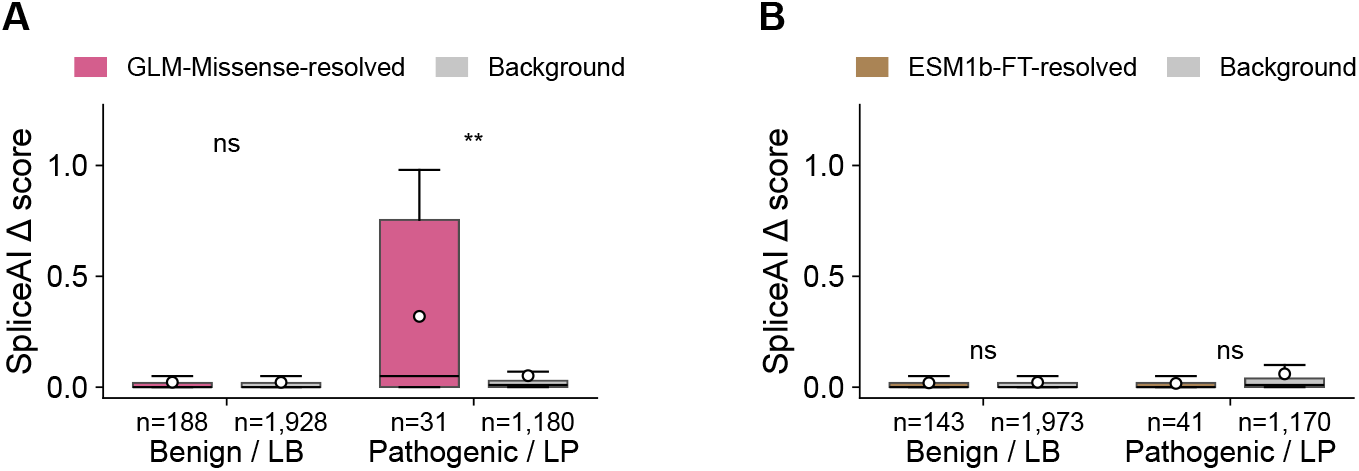
Pathogenic variants resolved by GLM-Missense, unlike those resolved by ESM1b-FT, are enriched for predicted splicing disruption. Maximum SpliceAI delta scores of resolved and background variants on the ClinVar later-release test set, shown separately for benign or likely benign and pathogenic or likely pathogenic variants. (A) GLM-Missense-resolved variants were correctly classified by GLM-Missense but misclassified by at least five of the seven established predictors (benign or likely benign *n* = 188, background *n* = 1,928; pathogenic or likely pathogenic *n* = 31, background *n* = 1,180). (B) ESM1b-FT-resolved variants, defined by the same criteria (benign or likely benign *n* = 143, background *n* = 1,973; pathogenic or likely pathogenic *n* = 41, background *n* = 1,170). Background variants are all other variants of the same class. Boxes represent the interquartile range with lines indicating medians, whiskers extending to the most extreme val-ues within 1.5 times the interquartile range, and white circles indicating means. Values beyond the whiskers are not shown. Two-sided Mann–Whitney *U*test: ns, not significant; ^∗∗^ *p <* 0.01.

**Fig. S14.**
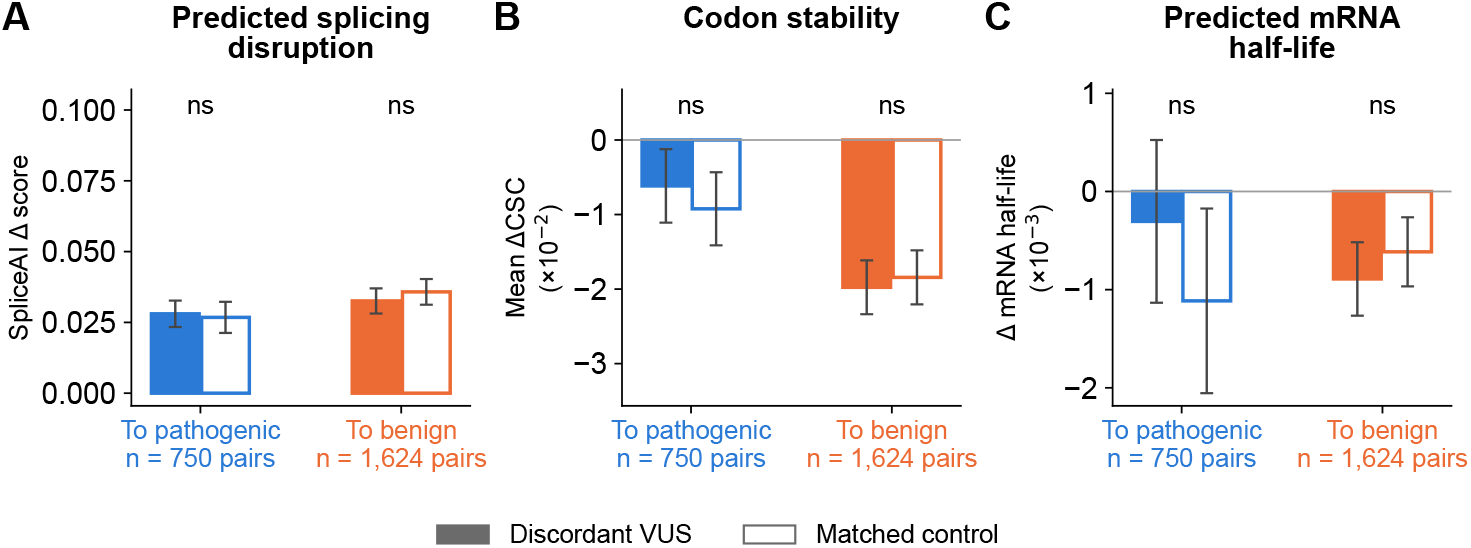
ESM1b-FT-discordant VUS show no significant differences from matched con-trols in splicing or mRNA-stability annotations. The analysis in Fig. 7 was repeated on the same held-out ClinVar missense VUS, with ESM1b-FT as the focal model. Discordant VUS (filled bars) were VUS for which ESM1b-FT made a confident prediction opposite to at least five of the seven comparator predictors. Matched controls (open bars) were selected using the same criteria as in Fig. 7. *To pathogenic* denotes discordant VUS that ESM1b-FT predicted to be pathogenic but at least five established predictors predicted to be benign (*n* = 750 pairs); *To benign* denotes the reverse (*n* = 1,624 pairs). (**A**) Maximum SpliceAI delta score. (**B**) Mean change in codon stability coefficient (ΔCSC). (**C**) Change in mRNA half-life predicted by Saluki. Bars represent means, and error bars indicate 95% confidence intervals of the means; y axes are the same as in Fig. 7. Two-sided Wilcoxon signed-rank tests on the pairs, with Benjamini–Hochberg adjustment over the six tests shown: ns, not significant.

**Table S1.** Five-fold cross-validation results for the top 10 LoRA configurations selected from the earlier-release strict validation set. The table reports cross-validation performance for the 10 highest-ranked LoRA models from Figure 3. Cross-validation was performed on the 6,250-variant subset of the earlier-release strict set using chromosome-exclusive folds. Models are ranked by the mean of AUROC and AUPRC ranks across folds.

| CV Rank | Val. Rank | Clf | Emb. | LoRA *r* | LR | Mean AUROC | STE | Mean AUPRC | STE |
|---|---|---|---|---|---|---|---|---|---|
| 1.0 | 1.0 | mlp | var-pos | 32 | 5e-5 | 0.850 | 0.006 | 0.819 | 0.006 |
| 2.5 | 2.0 | transformer | var-pos | 16 | 5e-5 | 0.847 | 0.005 | 0.814 | 0.005 |
| 2.5 | 4.0 | transformer | var-pos | 32 | 5e-5 | 0.847 | 0.005 | 0.814 | 0.003 |
| 4.0 | 9.5 | mlp | var-pos | 16 | 5e-5 | 0.845 | 0.005 | 0.811 | 0.008 |
| 6.0 | 3.0 | mlp | var-pos | 8 | 5e-5 | 0.844 | 0.005 | 0.807 | 0.006 |
| 6.0 | 7.5 | transformer | var-pos | 32 | 3e-5 | 0.843 | 0.005 | 0.808 | 0.007 |
| 6.0 | 9.0 | transformer | var-pos | 8 | 5e-5 | 0.843 | 0.004 | 0.808 | 0.008 |
| 8.0 | 6.5 | mlp | var-pos | 8 | 3e-5 | 0.842 | 0.006 | 0.804 | 0.009 |
| 9.5 | 6.5 | transformer | var-pos | 8 | 3e-5 | 0.838 | 0.005 | 0.801 | 0.009 |
| 9.5 | 7.0 | mlp | var-pos | 32 | 3e-5 | 0.839 | 0.006 | 0.799 | 0.010 |

**CV Rank:** mean of AUROC and AUPRC ranks from 5-fold cross-validation. **Val. Rank:** mean of AUROC and AUPRC ranks from the held-out earlier-release strict validation set. **Clf:** classifier head. **Emb.:** embedding; var-pos = variant-position embedding. **LoRA** ***r*****:** LoRA rank, controlling the dimensionality of the low-rank adaptation matrices. **LR:** learning rate used during fine-tuning. **Mean AUROC:** mean area under the ROC curve across folds. **Mean AUPRC:** mean area under the precision-recall curve across folds. **STE:** standard error of the mean across folds.

**Table S2.** Pathogenic or likely pathogenic ClinVar missense variants resolved by GLM-Missense. The table includes all pathogenic or likely pathogenic variants in the ClinVar later-release test set that were correctly classified by GLM-Missense but misclassified by at least five of the seven established predictors, together with SpliceAI scores, exon-exon junction distances, and ClinVar splicing-evidence status.

| ClinVar<br>ID | Chr | Pos | Ref<br>nt | Alt<br>nt | SpliceAI<br>score | Junction<br>dist | Splice<br>evidence |
|---|---|---|---|---|---|---|---|
| 4795219 | 3 | 14124672 | G | A | 0.09 | 18 |  |
| 4070964 | 3 | 25794304 | T | A | 0 | 213 |  |
| 4886898 | 3 | 184245333 | A | G | 0 | 26 |  |
| 4954121 | 2 | 26483490 | T | A | 0.01 | 29 |  |
| 4954913 | 2 | 47023465 | G | A | 0.48 | 1 |  |
| 4852563 | 2 | 70088650 | A | G | 0 | NA<sup>*a*</sup> |  |
| 8364 | 2 | 108753466 | C | T | 0 | 41 |  |
| 4876753 | 2 | 227094281 | C | G | 0.01 | 9 |  |
| 4082348 | 3 | 9434882 | G | A | 0.61 | 1 |  |
| 4974581 | 3 | 179234219 | A | T | 0 | 126 |  |
| 4819970 | 3 | 179234305 | G | A | 0 | 212 |  |
| 4796770 | 16 | 2094092 | C | G | 0.95 | 1 |  |
| 4875652 | 16 | 2135538 | C | G | 0 | 64 |  |
| 4928079 | 16 | 2176374 | G | C | 0 | 11 |  |
| 4928446 | 16 | 5078017 | G | A | 0.2 | 1 |  |
| 4762982 | 16 | 23634862 | C | G | 0.9 | 1 | Yes |
| 1713226 | 16 | 23641111 | T | G | 0.79 | 2 | Yes |
| 4755737 | 16 | 30964323 | G | T | 0.95 | 1 | Yes |
| 4686164 | 17 | 710564 | T | C | 0 | 32 |  |
| 4850612 | 17 | 3489255 | C | T | 0 | 21 |  |
| 4815395 | 17 | 3655101 | G | C | 0.9 | 1 |  |
| 4933576 | 17 | 8109321 | C | G | 0 | 23 |  |
| 2887278 | 17 | 44376320 | A | G | 0.01 | 13 |  |
| 4849901 | 17 | 50606984 | C | A | 0.05 | 6 |  |
| 4936238 | 17 | 59681695 | G | C | 0 | 49 |  |
| 4876164 | 17 | 61743013 | C | G | 0.98 | 1 | Yes |
| 4876516 | 17 | 80111025 | G | C | 0.83 | 1 |  |
| 4876515 | 17 | 80111983 | G | A | 0.08 | 1 |  |
| 4876673 | 17 | 80112100 | G | T | 0.47 | 1 |  |
| 431576 | 17 | 31200594 | A | G | 0.72 | 2 | Yes |
| 1745212 | 17 | 61849129 | C | G | 0.87 | 1 | Yes |

**ClinVar ID,** ClinVar variant accession identifier; **Chr,** chromosome; **Pos,** genomic position (hg38); **Ref nt,** reference nucleotide; **Alt nt,** alternate nucleotide; **SpliceAI score,** maximum SpliceAI delta score across acceptor gain, acceptor loss, donor gain, and donor loss; **Junction dist,** distance (nt) to the nearest exon-exon junction; **Splice evidence,** Yes if the ClinVar record includes splicing-relevant evidence.
<sup>*a*</sup>Variant lies in *PCBP1*, a single-exon gene with no exon-exon junction.

**Table S3.** Decision thresholds used to define resolved and discordant variants. For the seven established predictors, pathogenic and benign calls use the lower bound of the PP3 Supporting interval and the upper bound of the BP4 Supporting interval calibrated by ClinGen Bergquist et al. (2025); scores between the two thresholds receive no call. GLM-Missense and ESM1b-FT, which have no clinical calibration, are binarized at 0.5 in the resolved-variant analyses; in the analysis of discordant VUS, only confident calls (score *≥* 0.8 or *≤* 0.2) of these two models are used.

| Method | Pathogenic call | Benign call | Source |
|---|---|---|---|
| GLM-Missense | ≥ 0.5 | < 0.5 | this study |
| ESM1b-FT | ≥ 0.5 | < 0.5 | this study |
| AlphaMissense | ≥ 0.792 | ≤ 0.169 | Bergquist et al. (2025) |
| BayesDel (no AF) | ≥ 0.130 | ≤ −0.180 | Bergquist et al. (2025) |
| ESM1b | ≤ −10.7 | ≥ −6.3 | Bergquist et al. (2025) |
| MutPred2 | ≥ 0.737 | ≤ 0.391 | Bergquist et al. (2025) |
| REVEL | ≥ 0.644 | ≤ 0.290 | Bergquist et al. (2025) |
| VARITY_R | ≥ 0.675 | ≤ 0.251 | Bergquist et al. (2025) |
| VEST4 | ≥ 0.764 | ≤ 0.449 | Bergquist et al. (2025) |

## References

Adzhubei IA, Schmidt S, Peshkin L, et al (2010) A method and server for predicting damaging missense mutations. Nature Methods 7(4):248–249. 10.1038/nmeth0410-248

Agarwal V, Kelley DR (2022) The genetic and biochemical determinants of mRNA degradation rates in mammals. Genome Biology 23:245. 10.1186/s13059-022-02811-x

Bateman A, Martin MJ, Orchard S, et al (2023) UniProt: the Universal Protein Knowledgebase in 2023. Nucleic Acids Research 51(D1):D523–D531. 10.1093/nar/gkac1052

Benegas G, Ye C, Albors C, et al (2025) Genomic language models: opportunities and challenges. Trends in Genetics 41(4):286–302. 10.1016/j.tig.2024.11.013

Benjamini Y, Hochberg Y (1995) Controlling the false discovery rate: a practical and powerful approach to multiple testing. Journal of the Royal Statistical Society: Series B (Methodological) 57(1):289–300. 10.1111/j.2517-6161.1995.tb02031.x

Bergquist T, Stenton SL, Nadeau EAW, et al (2025) Calibration of additional computational tools expands ClinGen recommendation options for variant classification with PP3/BP4 criteria. Genetics in Medicine 27(6):101402. 10.1016/j.gim.2025.101402

Biderman D, Portes J, Gonzalez Ortiz JJ, et al (2024) LoRA learns less and forgets less. Transactions on Machine Learning Research URL https://openreview.net/forum?id=aloEru2qCG

Brandes N, Goldman G, Wang CH, et al (2023) Genome-wide prediction of disease variant effects with a deep protein language model. Nature Genetics 55:1512–1522. 10.1038/s41588-023-01465-0

Bueno-Martínez E, Sanoguera-Miralles L, Valenzuela-Palomo A, et al (2021) RAD51D aberrant splicing in breast cancer: identification of splicing regulatory elements and minigene-based evaluation of 53 DNA variants. Cancers 13(11):2845. 10.3390/cancers13112845

Capriotti E, Fariselli P (2022) Evaluating the relevance of sequence conservation in the prediction of pathogenic missense variants. Human Genetics 141(10):1649–1658. 10.1007/s00439-021-02419-4

Cartegni L, Chew SL, Krainer AR (2002) Listening to silence and understanding nonsense: exonic mutations that affect splicing. Nature Reviews Genetics 3(4):285–298. 10.1038/nrg775

Carter H, Douville C, Stenson PD, et al (2013) Identifying Mendelian disease genes with the variant effect scoring tool. BMC Genomics 14(Suppl 3):S3. 10.1186/1471-2164-14-S3-S3

Casadei S, Snyder MW, Woo I, et al (2026) Saturation genome editing reveals the functional impact of RAD51D and XRCC2 variants. bioRxiv 10.64898/2026.06.12.731983

Chen T, Guestrin C (2016) XGBoost: A scalable tree boosting system. In: Proceedings of the 22nd ACM SIGKDD International Conference on Knowledge Discovery and Data Mining, pp 785–794, 10.1145/2939672.2939785

Cheng J, Novati G, Pan J, et al (2023) Accurate proteome-wide missense variant effect prediction with AlphaMissense. Science 381(6664):eadg7492. 10.1126/science.adg7492

Christmas MJ, Kaplow IM, Genereux DP, et al (2023) Evolutionary constraint and innovation across hundreds of placental mammals. Science 380(6643):eabn3943. 10.1126/science.abn3943

Cooper GM, Shendure J (2011) Needles in stacks of needles: finding disease-causal variants in a wealth of genomic data. Nature Reviews Genetics 12(9):628–640. 10.1038/nrg3046

Curtis D (2025) Assessment of ability of a DNA language model to predict pathogenicity of rare coding variants. Journal of Human Genetics 70:603–607. 10.1038/s10038-025-01385-3

Dalla-Torre H, Gonzalez L, Mendoza-Revilla J, et al (2025) Nucleotide Transformer: building and evaluating robust foundation models for human genomics. Nature Methods 22:287–297. 10.1038/s41592-024-02523-z

Davydov EV, Goode DL, Sirota M, et al (2010) Identifying a high fraction of the human genome to be under selective constraint using GERP++. PLoS Computational Biology 6(12):e1001025. 10.1371/journal.pcbi.1001025

Dizkirici Tekpinar A, Tekpinar M (2025) A collaborative approach to improving missense mutational effect predictions in oncoproteins. medRxiv 10.1101/2025.05.15.25327738

Fairbrother WG, Yeh RF, Sharp PA, et al (2002) Predictive identification of exonic splicing enhancers in human genes. Science 297(5583):1007–1013. 10.1126/science.1073774

Feng BJ (2017) PERCH: a unified framework for disease gene prioritization. Human Mutation 38(3):243–251. 10.1002/humu.23158

Feng H, Wu L, Zhao B, et al (2025) Benchmarking DNA foundation models for genomic and genetic tasks. Nature Communications 16:10780. 10.1038/s41467-025-65823-8

Findlay GM, Boyle EA, Hause RJ, et al (2014) Saturation editing of genomic regions by multiplex homology-directed repair. Nature 513(7516):120–123. 10.1038/nature13695

Findlay GM, Daza RM, Martin B, et al (2018) Accurate classification of BRCA1 variants with saturation genome editing. Nature 562(7726):217–222. 10.1038/s41586-018-0461-z

Fowler DM, Fields S (2014) Deep mutational scanning: a new style of protein science. Nature Methods 11(8):801–807. 10.1038/nmeth.3027

Gerasimavicius L, Liu X, Marsh JA (2020) Identification of pathogenic missense mutations using protein stability predictors. Scientific Reports 10(1):15387. 10.1038/s41598-020-72404-w

Gerasimavicius L, Livesey BJ, Marsh JA (2022) Loss-of-function, gain-of-function and dominant-negative mutations have profoundly different effects on protein structure. Nature Communications 13(1):3895. 10.1038/s41467-022-31686-6

Grimm DG, Azencott CA, Aicheler F, et al (2015) The evaluation of tools used to predict the impact of missense variants is hindered by two types of circularity. Human Mutation 36(5):513–523. 10.1002/humu.22768

Hu EJ, Shen Y, Wallis P, et al (2022) LoRA: Low-rank adaptation of large language models. In: International Conference on Learning Representations, URL https://openreview.net/forum?id=nZeVKeeFYf9

Ioannidis NM, Rothstein JH, Pejaver V, et al (2016) REVEL: An ensemble method for predicting the pathogenicity of rare missense variants. American Journal of Human Genetics 99(4):877–885. 10.1016/j.ajhg.2016.08.016

Jaganathan K, Kyriazopoulou Panagiotopoulou S, McRae JF, et al (2019) Predicting splicing from primary sequence with deep learning. Cell 176(3):535–548.e24. 10.1016/j.cell.2018.12.015

Ji Y, Zhou Z, Liu H, et al (2021) DNABERT: pre-trained Bidirectional Encoder Representations from Transformers model for DNA-language in genome. Bioinformatics 37(15):2112–2120. 10.1093/bioinformatics/btab083

Kirkpatrick J, Pascanu R, Rabinowitz N, et al (2017) Overcoming catastrophic for-getting in neural networks. Proceedings of the National Academy of Sciences 114(13):3521–3526. 10.1073/pnas.1611835114

Landrum MJ, Lee JM, Benson M, et al (2018) ClinVar: improving access to variant interpretations and supporting evidence. Nucleic Acids Research 46(D1):D1062–D1067. 10.1093/nar/gkx1153

Landrum MJ, Chitipiralla S, Brown GR, et al (2020) ClinVar: improvements to accessing data. Nucleic Acids Research 48(D1):D835–D844. 10.1093/nar/gkz972

Lin Z, Akin H, Rao R, et al (2023) Evolutionary-scale prediction of atomic-level protein structure with a language model. Science 379(6637):1123–1130. 10.1126/science.ade2574

Liu X, Li C, Mou C, et al (2020) dbNSFP v4: a comprehensive database of transcript-specific functional predictions and annotations for human nonsynonymous and splice-site SNVs. Genome Medicine 12:103. 10.1186/s13073-020-00803-9

Liu Y, Ott M, Goyal N, et al (2019) RoBERTa: A robustly optimized BERT pretraining approach. arXiv preprint arXiv:190711692

Livesey BJ, Marsh JA (2020) Using deep mutational scanning to benchmark variant effect predictors and identify disease mutations. Molecular Systems Biology 16(7):e9380. 10.15252/msb.20199380

Livesey BJ, Marsh JA (2023) Updated benchmarking of variant effect predictors using deep mutational scanning. Molecular Systems Biology 19(8):e11474. 10.15252/msb.202211474

Majewski J, Schwartzentruber J, Lalonde E, et al (2011) What can exome sequencing do for you? Journal of Medical Genetics 48(9):580–589. 10.1136/jmedgenet-2011-100223

Maquat LE, Kinniburgh AJ, Rachmilewitz EA, et al (1981) Unstable *β*-globin mRNA in mRNA-deficient *β*^0^ thalassemia. Cell 27(3 Pt 2):543–553. 10.1016/0092-8674(81)90396-2

Marin FI, Teufel F, Horlacher M, et al (2024) BEND: Benchmarking DNA language models on biologically meaningful tasks. In: International Conference on Learning Representations

McCloskey M, Cohen NJ (1989) Catastrophic interference in connectionist networks: the sequential learning problem. Psychology of Learning and Motivation 24:109–165. 10.1016/S0079-7421(08)60536-8

McLaughlin HM, Ceyhan-Birsoy O, Christensen KD, et al (2014) A systematic approach to the reporting of medically relevant findings from whole genome sequencing. BMC Medical Genetics 15:134. 10.1186/s12881-014-0134-1

Meier J, Rao R, Verkuil R, et al (2021) Language models enable zero-shot prediction of the effects of mutations on protein function. In: Ranzato M, Beygelzimer A, Dauphin Y, et al (eds) Advances in Neural Information Processing Systems, vol 34. Curran Associates, Inc., pp 29287–29303, URL https://proceedings.neurips.cc/paper_files/paper/2021/file/f51338d736f95dd42427296047067694-Paper.pdf

Ng PC, Henikoff S (2003) SIFT: Predicting amino acid changes that affect protein function. Nucleic Acids Research 31(13):3812–3814. 10.1093/nar/gkg509

Ng PC, Henikoff S (2006) Predicting the effects of amino acid substitutions on protein function. Annual Review of Genomics and Human Genetics 7:61–80. 10.1146/annurev.genom.7.080505.115630

Nguyen E, Poli M, Faizi M, et al (2023) HyenaDNA: Long-range genomic sequence modeling at single nucleotide resolution. In: Advances in Neural Information Processing Systems, URL https://proceedings.neurips.cc/paper_files/paper/2023/hash/86ab6927ee4ae9bde4247793c46797c7-Abstract-Conference.html

Pejaver V, Urresti J, Lugo-Martinez J, et al (2020) Inferring the molecular and phenotypic impact of amino acid variants with MutPred2. Nature Communications 11:5918. 10.1038/s41467-020-19669-x

Pejaver V, Byrne AB, Feng BJ, et al (2022) Calibration of computational tools for missense variant pathogenicity classification and ClinGen recommendations for PP3/BP4 criteria. American Journal of Human Genetics 109(12):2163–2177. 10.1016/j.ajhg.2022.10.013

Pollard KS, Hubisz MJ, Rosenbloom KR, et al (2010) Detection of nonneutral sub-stitution rates on mammalian phylogenies. Genome Research 20(1):110–121. 10.1101/gr.097857.109

Presnyak V, Alhusaini N, Chen YH, et al (2015) Codon optimality is a major determinant of mRNA stability. Cell 160(6):1111–1124. 10.1016/j.cell.2015.02.029

Rastogi R, Chung R, Li S, et al (2025) Critical assessment of missense variant effect predictors on disease-relevant variant data. Human Genetics 144(2–3):281–293. 10.1007/s00439-025-02732-2

Rehm HL, Berg JS, Brooks LD, et al (2015) ClinGen—the Clinical Genome Resource. New England Journal of Medicine 372(23):2235–2242. 10.1056/NEJMsr1406261

Richards S, Aziz N, Bale S, et al (2015) Standards and guidelines for the interpretation of sequence variants: a joint consensus recommendation of the American College of Medical Genetics and Genomics and the Association for Molecular Pathology. Genetics in Medicine 17(5):405–424. 10.1038/gim.2015.30

Rives A, Meier J, Sercu T, et al (2021) Biological structure and function emerge from scaling unsupervised learning to 250 million protein sequences. Proceedings of the National Academy of Sciences 118(15):e2016239118. 10.1073/pnas.2016239118

Sabarinathan R, Tafer H, Seemann SE, et al (2013) RNAsnp: Efficient detection of local RNA secondary structure changes induced by SNPs. Human Mutation 34(4):546–556. 10.1002/humu.22273

Sauna ZE, Kimchi-Sarfaty C (2011) Understanding the contribution of synonymous mutations to human disease. Nature Reviews Genetics 12(10):683–691. 10.1038/nrg3051

Schiff Y, Kao CH, Gokaslan A, et al (2024) Caduceus: Bi-directional equivariant long-range DNA sequence modeling. In: Proceedings of the 41st International Conference on Machine Learning, Proceedings of Machine Learning Research, vol 235. PMLR, pp 43632–43648, URL https://proceedings.mlr.press/v235/schiff24a.html

Soemedi R, Cygan KJ, Rhine CL, et al (2017) Pathogenic variants that alter protein code often disrupt splicing. Nature Genetics 49:848–855. 10.1038/ng.3837

Starita LM, Ahituv N, Dunham MJ, et al (2017) Variant interpretation: Functional assays to the rescue. American Journal of Human Genetics 101(3):315–325. 10.1016/j.ajhg.2017.07.014

Stefl S, Nishi H, Petukh M, et al (2013) Molecular mechanisms of disease-causing missense mutations. Journal of Molecular Biology 425(21):3919–3936. 10.1016/j.jmb.2013.07.014

Stergachis AB, Haugen E, Shafer A, et al (2013) Exonic transcription factor binding directs codon choice and affects protein evolution. Science 342(6164):1367–1372. 10.1126/science.1243490

Tavtigian SV, Harrison SM, Boucher KM, et al (2020) Fitting a naturally scaled point system to the ACMG/AMP variant classification guidelines. Human Mutation 41(10):1734–1737. 10.1002/humu.24088

Tennessen JA, Bigham AW, O’Connor TD, et al (2012) Evolution and functional impact of rare coding variation from deep sequencing of human exomes. Science 337(6090):64–69. 10.1126/science.1219240

Veiner M, Supek F (2026) The DNA dialect: a comprehensive guide to pretrained genomic language models. Molecular Systems Biology 22:309–332. 10.1038/s44320-025-00184-4

Walker LC, de la Hoya M, Wiggins GAR, et al (2023) Using the ACMG/AMP frame-work to capture evidence related to predicted and observed impact on splicing: Recommendations from the ClinGen SVI Splicing Subgroup. American Journal of Human Genetics 110(7):1046–1067. 10.1016/j.ajhg.2023.06.002

Wang Y, Cai Z, Zeng Q, et al (2025) Genomic Touchstone: benchmarking genomic language models in the context of the central dogma. bioRxiv 10.1101/2025.06.25.661622

Weile J, Roth FP (2018) Multiplexed assays of variant effects contribute to a growing genotype–phenotype atlas. Human Genetics 137(9):665–678. 10.1007/s00439-018-1916-x

Whalen S, Schreiber J, Noble WS, et al (2022) Navigating the pitfalls of applying machine learning in genomics. Nature Reviews Genetics 23(3):169–181. 10.1038/s41576-021-00434-9

Wu Q, Medina SG, Kushawah G, et al (2019) Translation affects mRNA stability in a codon-dependent manner in human cells. eLife 8:e45396. 10.7554/eLife.45396

Wu Y, Liu H, Li R, et al (2021) Improved pathogenicity prediction for rare human missense variants. American Journal of Human Genetics 108(10):1891–1906. 10.1016/j.ajhg.2021.08.012

Xiong HY, Alipanahi B, Lee LJ, et al (2015) RNA splicing. The human splicing code reveals new insights into the genetic determinants of disease. Science 347(6218):1254806. 10.1126/science.1254806

Zhou Z, Ji Y, Li W, et al (2024) DNABERT-2: Efficient foundation model and benchmark for multi-species genomes. In: International Conference on Learning Representations

